# Decoding morphogen patterning of human neural organoids with a multiplexed single-cell transcriptomic screen

**DOI:** 10.1101/2024.02.08.579413

**Authors:** Fátima Sanchís-Calleja, Akanksha Jain, Zhisong He, Ryoko Okamoto, Charlotte Rusimbi, Pedro Rifes, Gaurav Singh Rathore, Malgorzata Santel, Jasper Janssens, Makiko Seimiya, Jonas Simon Fleck, Agnete Kirkeby, J. Gray Camp, Barbara Treutlein

## Abstract

Morphogens, secreted signalling molecules that direct cell fate and tissue development, are used to direct neuroepithelial progenitors towards discrete regional identities across the central nervous system. Neural tissues derived from pluripotent stem cells in vitro (neural organoids) provide new models for studying neural regionalization, however, we lack a comprehensive survey of how the developing human neuroepithelium responds to morphogen cues. Here, we produce a detailed map of morphogen-induced effects on the axial and regional specification of human neural organoids using a multiplexed single-cell transcriptomics screen. We find that the timing, concentration, and combination of morphogens strongly influence organoid cell type and regional composition, and that cell line and neural induction method strongly impact the response to a given morphogen condition. We apply concentration gradients in microfluidic chips or a range of static concentrations in multi-well plates to explore how human neuroepithelium interprets morphogen concentrations and observe similar dose-dependent induction of patterned domains in both scenarios. Altogether, we provide a detailed resource that supports the development of new regionalized neural organoid protocols and enhances our understanding of human central nervous system patterning.

## Introduction

The elaborate architecture and functionality of the human central nervous system (CNS) arises from tightly coordinated patterning events initiated in early stages of development when the neural plate forms and gives rise to the neural tube. Throughout CNS development, a limited set of morphogens with unique activity patterns interact in space and time to produce a diverse range of regions and cell types. Morphogens are secreted from distinct locations in the developing neuroepithelium and govern neural regionalization in part by establishing gradients that instruct compartmentalization into multiple domains on the anteroposterior and dorso-ventral axes ^1–3^. Current knowledge of CNS regionalization mostly originates from studies using non-human animal models with divergent cytoarchitecture, cell behavior and gene expression patterns compared to humans ^4–7^. Research on human tissue has been hampered by largely unattainable primary samples and ethical considerations, however the advent of stem cell-based neural differentiation methods has enabled the study of human-specific developmental features in vitro ^6–12^. While monolayer neural cultures present basic rosette-like arrangements, neural organoids provide a three-dimensional environment where cells self-organize cytoarchitectural and morphogenetic features with similarities to the primary human tissue ^13,14^. Human stem cell-derived neuroepithelium presents a unique opportunity to systematically study how morphogen timing, concentration, and combination impacts a naive neuroepithelium.

Numerous approaches have been developed to generate reproducibly patterned neural organoids, such as organoids with intrinsic inducible organizers ^15^, self-folding systems with dorso-ventral patterning ^16^, and a wide range of protocols for the generation of regionalized neural organoids of forebrain, midbrain, hindbrain or retinal-like tissue fates ^17–22^. Although these advances have increased our understanding of neural tissue regionalization and consolidated neural organoids as a leading model system for neurodevelopmental studies ^13,23^, it is still challenging to integrate the findings obtained in each study and evaluate their relevance for human-specific biology. Different guided organoid protocols aimed towards the same brain region can produce different outcomes, as they employ different culture settings that might affect tissue architecture, regional patterning, cell type diversity and maturation rates. In addition, the development of new protocols is still inefficient, since the assessment of different culture conditions most often relies on bulk measurements of a few marker genes with low organoid throughput.

To unify our understanding of early neural regionalization, we implement a systematic approach to assess morphogen effects on developing human neural organoids. We supply patterning molecules at multiple time points, concentrations, and combinations, and use single cell-RNA sequencing with cell hashing-based sample multiplexing ^24^ to assess neural progenitor regionalization. We show that the delivery of SHH, WNT, FGF8, RA, BMP4 and BMP7 pathway modulators in short pulses over different time windows induces strong changes in organoid composition and increasing concentration steps in enlarged treatment windows modulate the proportion of each regional identity within the same tissue. Guided by this analysis, we use combinations of morphogens to steer axial and regional fates in a more precise manner. Our experiments also revealed significant variability across human pluripotent stem cell (hPSC) lines and neural induction methods. We demonstrate that the generated organoids covered a substantial portion of the neural tube domains observed in vivo, with each protocol producing a composite of brain regional identities. Finally, we show that a SHH morphogen gradient and discontinuous SHH morphogen steps similarly recapitulate dorso-ventral patterning of the human forebrain. Altogether, our work provides an in-depth exploration of morphogen-induced patterning of human neural organoids, contributing to the understanding of human neurodevelopment and serving as a reference for the accelerated development of optimized organoid models.

## Results

### A single-cell atlas of human neural organoid patterning

We established an experimental workflow to systematically test the timing, concentration, and combinatorics of patterning molecules using cell hashing-based multiplexed single-cell RNA-sequencing. Neural organoids were generated in 96 well plates, in which aggregated multipotent progenitor cells (embryoid bodies, EBs) were cultured in neural induction media to generate a naive, unguided neuroepithelium (Fig. 1A). Developing organoids were then subjected to a panel of morphogen treatments with varying timing, concentration, and combination of the molecules CHIR99021 (CHIR, WNT pathway activator), XAV939 (WNT pathway inhibitor), rhSHH/CS24II (SHH) and Purmorphamine (PM), FGF-8B (FGF8), Retinoic Acid (RA), rhBMP-4 (BMP4), rhBMP-7 (BMP7) and Cyclopamine (Cyc) (Figures S1 and S2, Supplementary Tables S1, S2). On day 21, multiple neural organoids of each condition were pooled and dissociated into single-cell suspensions, followed by cell hashing with barcoded antibodies ^24^ and processing for single-cell transcriptome sequencing (3’ mRNA, 10X Genomics) (Fig. 1B). Altogether, we collected 100,538 cells including cells from untreated organoids (one control condition per 96-well plate or organoid batch) and 97 morphogen treatment conditions.

**Fig. 1.**
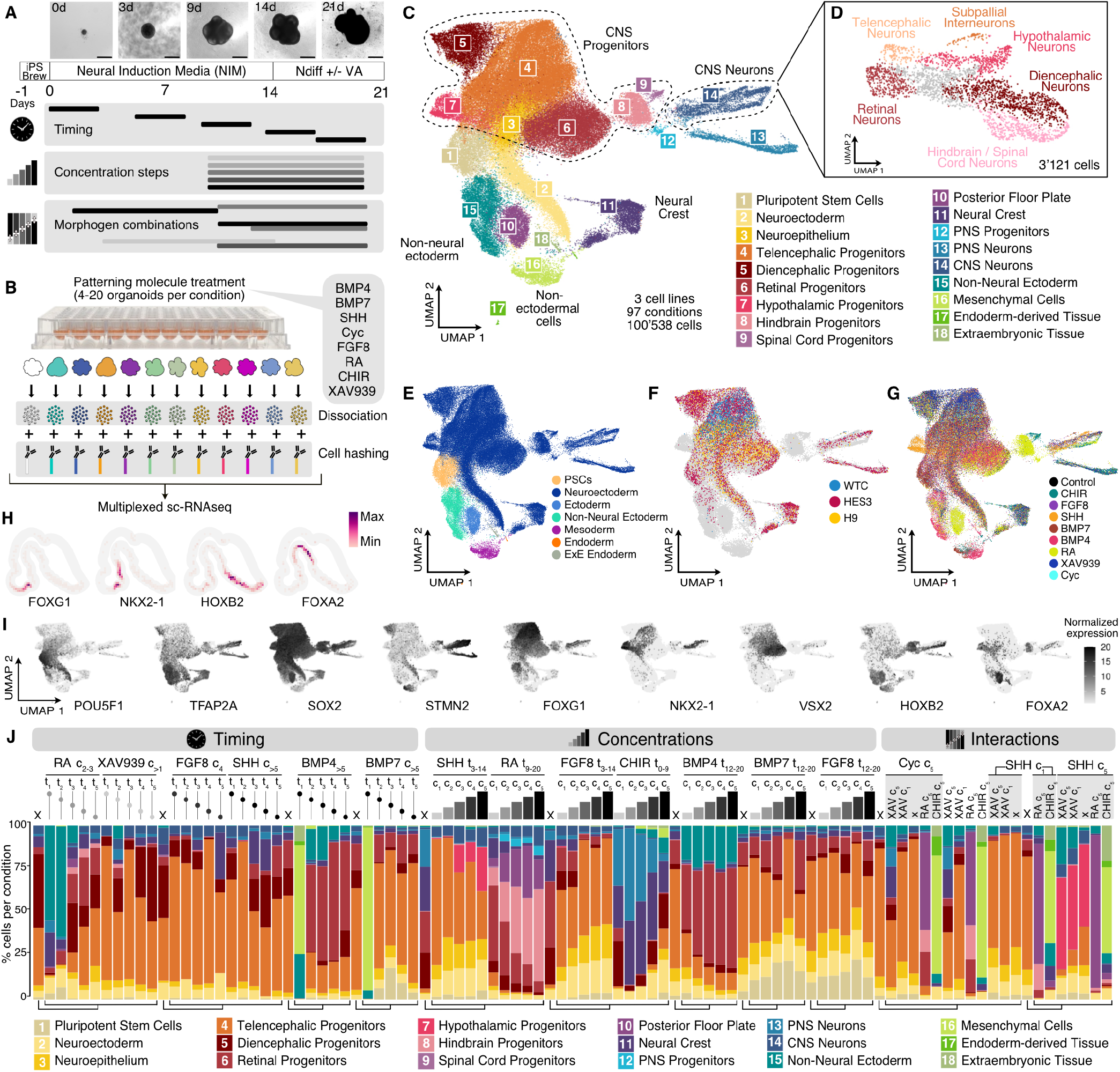
A single-cell transcriptomic atlas of human neural organoid patterning. (A) Experimental timeline showing morphogen treatment modalities in this study, an overview of the organoid culture protocol employed and representative brightfield images at different developmental stages (cell line = WTC). Scale bar = 500 µm. (B) Overview of the experimental approach to obtain multiplexed sc-RNAseq readouts from treated organoids. (C) RSS-based UMAP projection of all cells in the dataset, colored by cluster identity. (D) RSS based-UMAP projection of subclustered CNS Neurons, showing different regional identities in the same colors as their corresponding progenitors shown in (C). (E) Whole dataset RSS-UMAP projection colored by germ layer, showing off-target cells mostly generated by morphogen treatments. (F) RSS-UMAP projection highlighting untreated (control) cells from each cell line of origin. Cells from morphogen treatment conditions are shown behind in gray. (G) Same projection highlighting cells treated with a single morphogen and control cells. Cells from combined morphogen treatments are colored in gray in the background. (H) Voxel maps showing the normalized expression of regional marker genes in the developing human brain dataset published in Braun et al, 2023 ^26^. (I) Feature plots for representative marker genes (POU5F1, stem cells; TFAP2A, non-neural ectoderm or hindbrain; SOX2, neural stem cells; STMN2, neurons; FOXG1, telencephalon; NKX2-1, ventral telencephalon; VSX2, retina; HOXB2, hindbrain; FOXA2, floor plate). (J) Bar plots showing the cell type composition of each condition for HES3-derived organoids exposed to single time point, concentration steps and an example subset of combinatorial morphogen treatments. Concentrations are indicated by c1-5 and treatment windows by t0-21, indicating the start and end of morphogen exposure in days. Lines with dots represent timing experiments (single morphogen pulse), the dot representing the time-frame when the treatment took place (up is early, down is late). Concentration experiments are represented by progressively taller bars. The shading of the dots and bars in timing and concentration experiments represents the concentration used (darker shading means higher concentration). “X” indicates untreated (control) organoids. Correspondences between controls and experimental conditions are indicated by the tree below the bar plots. On the right, combinatorial experiments (“Interactions”) are labeled with the antero-posterior morphogen in vertical and the dorso-ventral morphogen in horizontal; for simplicity only concentrations are indicated (refer to Fig. S1 for timing information) and FGF8 combinations are not shown.

We integrated the data using Reference Similarity Spectrum (RSS) ^25^ comparison to an early developing human brain cell atlas reference ^26^ and embedded the dataset in 2D with Uniform Manifold Approximation and Projection (UMAP) ^27^ (Fig. 1C). Analysis of cluster marker gene expression (Fig. S3) revealed cell identities including diverse CNS progenitors and neurons of telencephalon, diencephalon, retina, hypothalamus, hindbrain, and spinal cord domains, as well as neural crest, non-neural ectoderm, mesoderm, endoderm and extraembryonic tissue (Fig. 1E). Further sub-clustering of CNS neurons revealed different regional identities consistent with the identities observed in CNS progenitors (Fig. 1D). We found that organoid cells from all PSC lines predominantly localized to neural clusters in the absence of any morphogen treatment (Fig. 1F), whereas treated organoids generated distinct clusters that resembled neural regions absent in control organoids, such as hindbrain and hypothalamus as well as other non-neural tissues (Fig. 1G-I).

Our single-cell resolved RNAseq readout allowed us to quantify changes in organoid cell type composition for each specific morphogen treatment (Fig. 1J). Few conditions showed homogenous cell type composition, while most conditions displayed an array of cellular identities spanning differentiation states between progenitors and neurons patterned to different brain regions. Interestingly, higher morphogen concentrations did not promote more homogenous cell type composition in the organoids, but instead generated new cell types that gradually replaced the initial identities. Altogether, we provide a single-cell transcriptome reference of human neural organoid patterning to explore principles of cell fate acquisition in response to extrinsic morphogen exposure.

### Effect of morphogen timing and concentration on regional cell identities in developing neural organoids

To identify effects of each morphogen at the cell type level, we calculated cluster enrichment/depletion scores for each treatment (Fig. 2A-B, Fig. S4). Most morphogen pathway manipulations resulted in enrichment or depletion of one or more clusters. Longer morphogen treatments (morphogen concentration experiment, Fig. 2B) induced the most significant changes in cell type composition. However, single pulses of certain treatments also showed remarkable effects, such as Retinoic Acid promoting non-neural ectoderm at early time points but retinal and hindbrain progenitors when supplied at later time points (Fig. 2A and Fig. 2C, Timing panel). Interestingly, late and long exposure (day 9-20) to the dose employed in the timing experiment (c2-3) promoted retinal progenitor production over hindbrain, whereas higher doses progressively enriched for hindbrain and posterior floor plate (Fig. 2C, Concentrations panel). In situ Hybridization Chain Reaction (HCR) stainings confirmed the presence of more retinal progenitors (SFRP2+) at lower RA concentrations and more ventralized and posterior fates (FOXA2+) at higher RA concentrations (Fig. 2D).

**Fig. 2.**
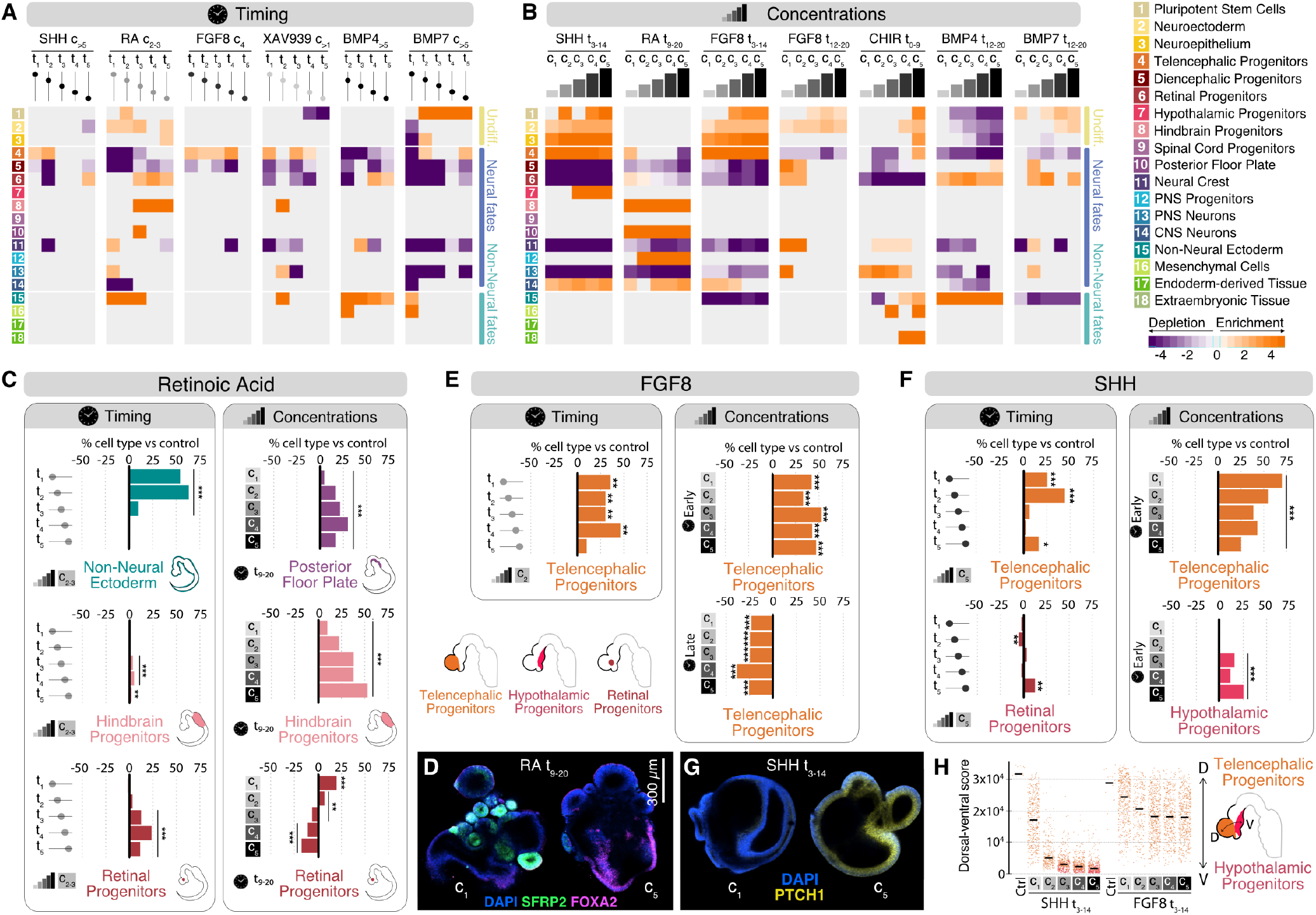
Morphogen timing and concentrations determine regional cell identities in developing neural organoids. (A,B) Heatmaps showing enrichment/depletion scores for timing (A) and concentration (B) experiments in HES3-derived organoids. The shading of dots and bars on top of the heatmap indicates the employed dose and the dots along a line represent the time point of patterning molecule addition. (C) Effects of different RA treatment windows (left) and concentration steps (right) on the abundance of retinal and hindbrain progenitors, non-neural ectoderm and posterior floor plate in 21-day-old neural organoids. (D) HCR in situ hybridization for SFRP2 (retinal marker gene) and FOXA2 comparing RA treatment conditions c1 and c5. (E) Effects of different FGF8 treatment windows (left) and concentration steps (right) at early (day 3-14) and late (day 12-20) time windows on the abundance of telencephalic progenitor populations at 21 days. Bottom left: Spatial representation of the clusters enriched by FGF8 and SHH in a schematic representation of a 4 GW human embryo. (F) Effects of different SHH treatment windows (left) and concentration steps (right) on the abundance of retinal, telencephalic and hypothalamic progenitor populations in 21 day-old neural organoids. (G) HCR in situ hybridization for PTCH1 (hypothalamic marker gene and target gene of SHH signaling) comparing SHH treatment conditions c1 and c5. (H) Dorso-ventral axis score values for different concentration steps of SHH and FGF8 with their corresponding controls showing higher ventralization in SHH-treated organoids, which reach the hypothalamic compartment. *** p-value < 0.001, ** 0.001< p-value < 0.01, * 0.01 < p-value < 0.05 (Bonferroni adjusted).

SHH and FGF8 both induced telencephalic fates when supplied in short pulses (timing experiments, Fig. 2E-F, Timing panels), with the SHH patterning window being more restricted to earlier time points (Fig. 2A and Fig. 2F, Timing panel) while FGF8 maintained this activity over a longer time window (Fig. 2A and Fig. 2E, Timing panel). However, when exposing the organoids to FGF8 for a longer time (concentration experiments), we observed depletion of telencephalic fates at later time points (day 12-20), in contrast to robust enrichment in early timepoints (day 3-14) (Fig. 2B and 2E, Concentrations panel). In the case of SHH, late exposure to a single pulse favored the production of retinal progenitors together with a slight enrichment in telencephalic identities (Fig. 2F, Timing panel). Progressively higher concentrations induced a transition from telencephalic to hypothalamic progenitors (Fig. 2F, Timing panel; Fig. 2G), consistent with the spatial concentration gradient of SHH in vivo, which is highest at the ventral most part of the developing neural tube from where it diffuses ^28–30^.

Since we observed similar levels of telencephalic progenitor enrichment in FGF8 and SHH treatments, we next compared their influence on dorso-ventral (DV) and antero-posterior (AP) patterning within the telencephalon. To this aim, we calculated DV and AP scores for each cell using marker genes varying along these axes (Fig. 2H and Fig. S5A). We observed that SHH exposure had a stronger anteriorizing effect on telencephalic progenitor cells than FGF8 (Fig. S5A-B). This finding is surprising since it has been shown in the mouse that FGF8 is secreted from the Anterior Neural Ridge (ANR) to promote anteriorization of the developing telencephalon, potentially suggesting a differential role of FGF8 in mouse and human brain patterning ^31^. With regard to the DV axis, SHH had stronger ventralizing effects than FGF8, generating hypothalamic floor plate cells at the highest concentrations (Fig. 2F and 2H).

The BMP family of ligands is known for partial redundancy and overlapping expression of its members, leading to compensation of deficits in expression of a given ligand ^32–36^. However, we still lack a unified dissection of their patterning roles. To better understand the logic of BMP-induced patterning, BMP4 and BMP7 were tested in equimolar steps to compare their effects on developing neural organoids. In the timing experiments, both BMP4 and BMP7 prevented neural specification when supplied in early time windows, consistent with their role in vivo ^37^ (Fig. 2A). While BMP4 treatment on days 0-3 caused disintegration of organoids, BMP7 induced a nearly 90% enrichment in mesenchymal cells (Fig. S4J). After day 3, BMP4 induced NNE and progressively guided towards retina cell fates at later time points (Fig. 2A and Fig. S4I). Instead, BMP7 guided cells towards pluripotency, even when added at later time points (Fig. S4J).

When supplying BMP4 over an extended period of time (days 12-20), increase in concentration led to consistent increase in the percentage of retinal progenitors, while the proportion of NNE cells remained unchanged (Fig. 2B, Fig. S4L). BMP7 also showed a dose-dependent induction of retinal progenitor fate, though less efficiently than BMP4 (Fig. 2B, Fig. S4M). We calculated differential expression between control, BMP4 and BMP7 conditions (Fig. S5D-E) and found very similar expression patterns in control and BMP7 treated organoids, whereas BMP4 treated cells activated MSX2 and TBX genes, implicated in neural crest development and dorsal neural tube formation ^38,39^. BMP4 also activated the expression of FOXE3 and SAMD11, which are essential for optic cup morphogenesis ^40,41^, consistent with the promotion of retinal fates.

Inhibition of the WNT pathway (XAV939) at early and mid time points (days 0-3, 10-13, and 14-17) led to a significant enrichment in telencephalic progenitors (Fig. 2A, Fig. S4C). Activation of the WNT pathway via exposure to CHIR caused strong dorsalization and posteriorization, as shown by the production of neural crest fates and PNS neurons (Fig. 2B, Fig. S4F). At highest concentrations, CHIR additionally guided cells towards non-neural identities such as mesenchyme (Fig. 2B).

### Morphogen conditions differentially activate transcription factor regulons in developing neural organoids

To explore how morphogens differentially activate transcription factors (TFs) and their downstream targets (regulons), we performed a gene regulatory network inference analysis using SCENIC ^42^ and linked morphogen conditions with regulon activities (Fig. 3A). Each morphogen treatment associated with a specific set of regulons, which showed enriched activity in a particular cell cluster. Notably, some regulons were associated with two or more morphogens, apparent as interconnected nodes in the network graph, such as regulons linked to both SHH and XAV939 (e.g. TCF7L2, NKX2-1). We next examined the dependency of regulon activity on morphogen timing and concentration, finding both positive and negative correlations (Fig. 3B-C and Fig. S6, S8). Retinoic Acid regulons such as HOXC4, POU3F2 and FOXA2 were dependent on both timing and concentration (Fig. 3D and Fig. S7, S8), whereas BMP4 regulons were highly dependent on timing, but not concentration (Fig. 3E and Fig. S7, S8). Inhibition of the WNT pathway with XAV939 mildly modulated regulon activity in a time-dependent manner (Fig. 3B and Fig. S7), whereas WNT pathway activation using CHIR showed a clear dependency on morphogen concentration. Interestingly, although the activity of some posteriorizing regulons like PAX7 was positively correlated to CHIR concentration, most CHIR-associated regulons were related to neurogenesis and showed negative correlation to CHIR concentration, indicating a possible suppression of neural fates under our culture conditions (Fig. 3C and Fig. S8).

**Fig. 3.**
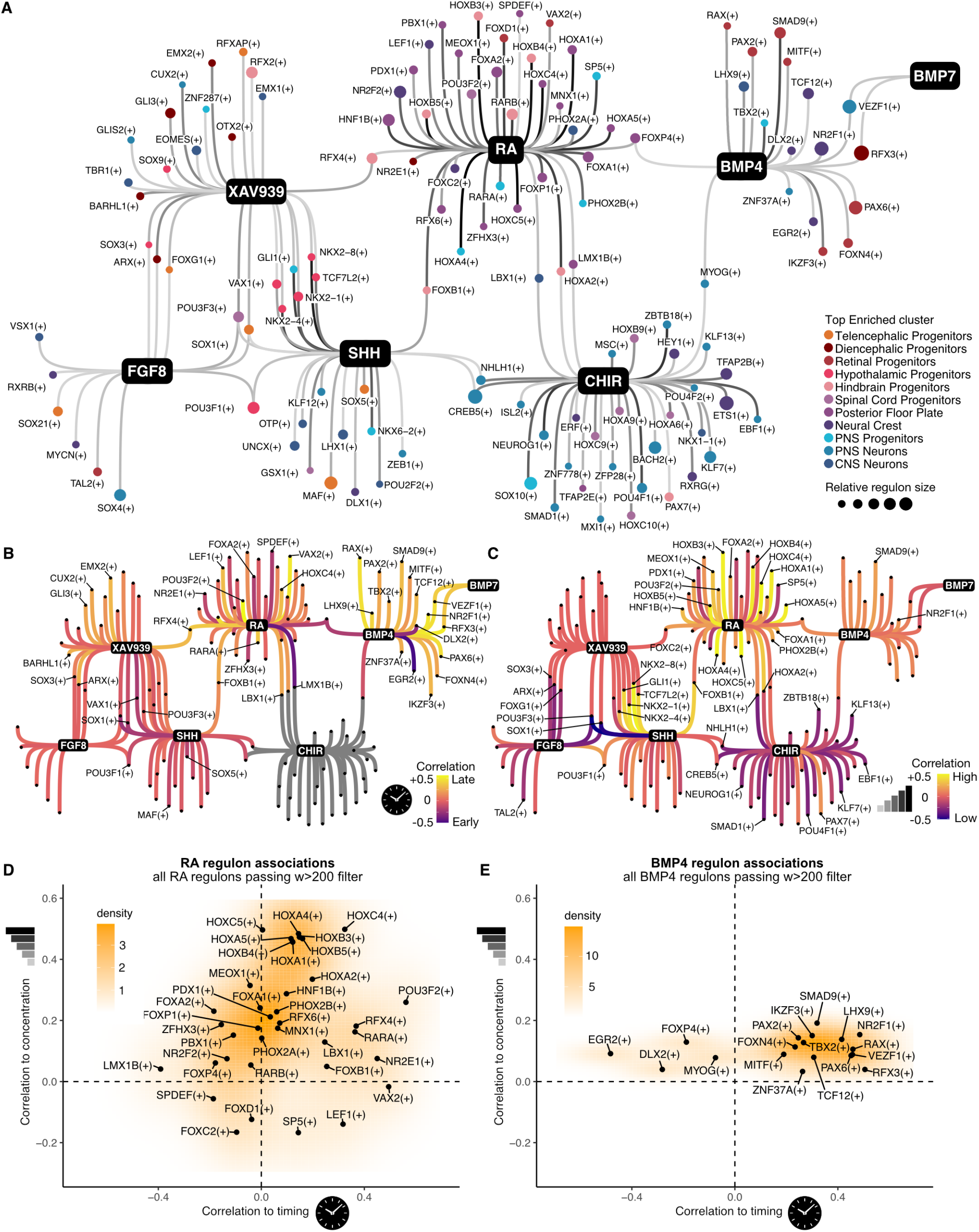
Morphogen conditions differentially activate transcription factor regulons in developing neural organoids. (A) Morphogen-guided regulon network for all tested patterning molecules. Each black node represents a morphogen that is connected to associated regulons (showing correlating activity to morphogen treatment). The size of each regulon node indicates the relative size of the regulon and the color refers to the cluster where that regulon registers maximum activity (excluding non-neural and undifferentiated clusters). (B,C) Same regulon network with edges colored according to the correlation to timing (B) and concentration (C). Positive correlation to timing means activity at later time points, whereas negative correlation implies early patterning window. For concentrations, negative correlation indicates regulons induced at low doses, while positive correlation points to dose-dependent increases in regulon activity. (D) Scatterplot of RA-associated regulon correlation to timing and concentration, showing wide dependency on these variables. (E) Scatterplot of BMP4-associated regulon correlation to timing and concentration, showing low dependency on concentration and high dependency on timing.

SHH- and FGF8-linked regulons did not correlate strongly with timing but showed strong dependence on concentration (Fig. 3B-C and Fig. S6). Although both morphogens shared associations with POU3F1(+), POU3F3(+) and SOX1(+) regulons, changes in SHH concentration generally led to stronger effects than changes in FGF8 concentration (Fig. S6 and S8). In addition, NKX2.1(+), POU3F1(+) and GLI1(+) regulon activation were dependent on SHH concentration, while POU3F1(+) and VSX1(+) regulon activation correlated with FGF8 concentration (Fig. S6 and S8). BMP7, like BMP4, showed higher dependency on timing than concentrations. However, it showed much weaker effects, with only 2 regulons passing our threshold for significant associations to BMP7 (weight > 200): VEZF1(+) and RFX3(+), also linked to BMP4 (Fig. 3B-C, Fig. S6-S8).

### A combination panel to probe morphogen interactions

To probe relevant morphogen combinations, we selected timing and concentration conditions and designed a combination scheme to reach most domains of the CNS (Fig. 4A). XAV939, RA and CHIR were used to pattern along the anteroposterior axis, whereas cyclopamine and SHH were applied to modulate along the dorsoventral axis. We designed additional conditions including FGF8 together with other morphogens that may generate ANR or midbrain-hindbrain boundary (MHB) domains (Fig. 4A, purple boxes).

**Fig. 4.**
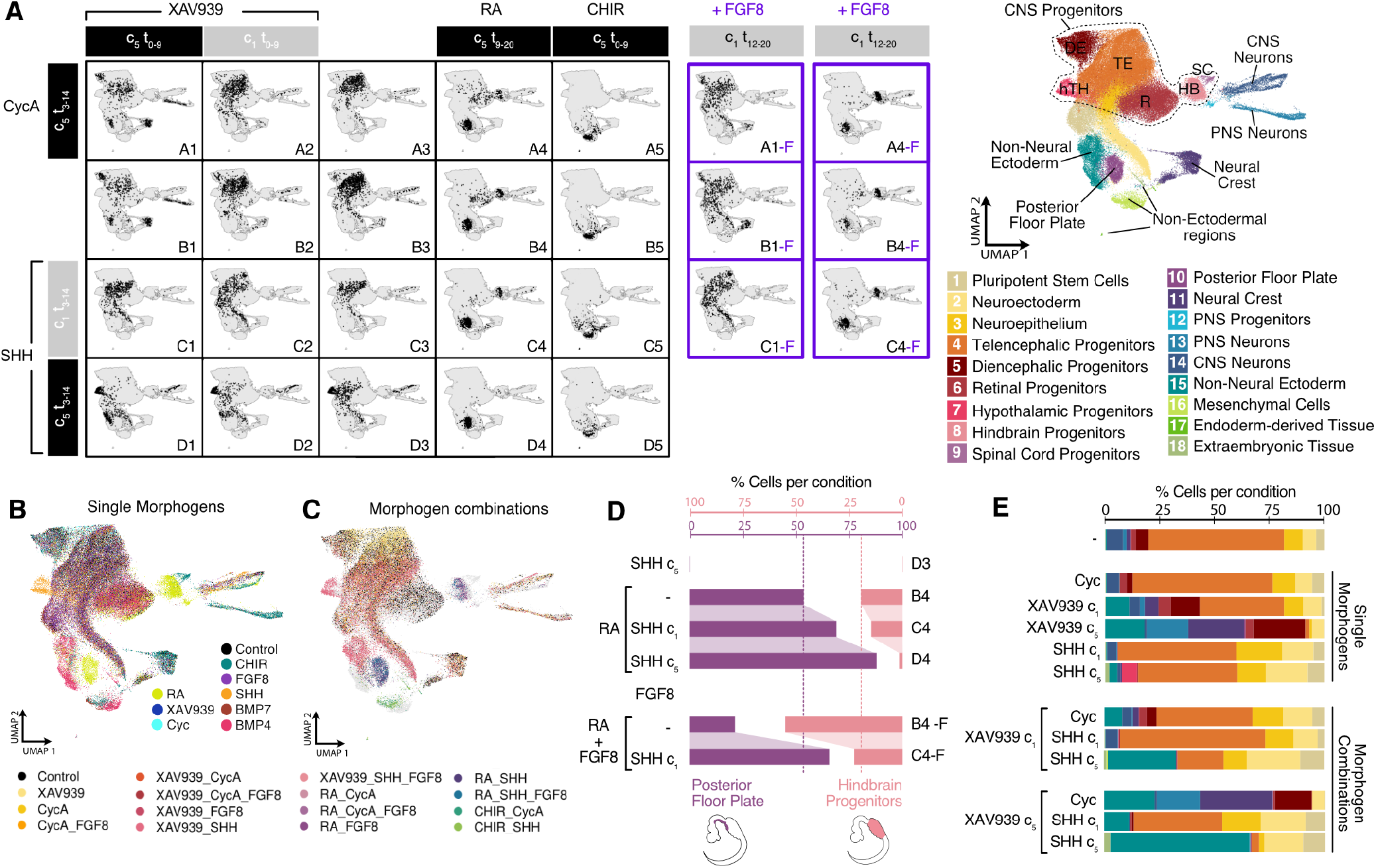
Morphogen combinations regulate neural cell fates. (A) Left, RSS-UMAP projections highlighting the cells corresponding to each coordinate of the morphogen treatment panel. Middle, in purple, treatments with the addition of FGF8 on top of the corresponding morphogens. A reference UMAP with cell type annotations is shown on the right. (B,C) RSS-UMAP projections showing cells from organoids treated with one single morphogen (B) or a combinatorial treatment (C). (D) Enrichment of hindbrain vs posterior floor plate in different conditions under RA, SHH and FGF8 treatment. The basal levels of posterior floor plate and hindbrain progenitors under treatment with only RA are shown with a discontinued line in purple or pink, respectively. (E) Cell type composition of combinatorial experiments which involve treatment with XAV939, SHH and cyclopamine (Cyc), showing the interplay between these morphogens.

Orthogonal combinations of patterning molecules did not generate new regional identities in treated organoids (Fig. 4A-C). Instead, some of the morphogen combinations created organoids enriched for specific regional identities, while others generated organoids with a complex combination of different identities (Fig. 4D-E and Fig. S2). For example, we found that the balance between hindbrain progenitors and posterior floor plate is regulated by RA, FGF8 and SHH (Fig. 4D). Organoids treated with RA alone generated a majority of posterior floor plate (53%) and a smaller proportion of hind-brain progenitors (19.5%). Higher SHH concentrations combined with RA increased the proportion of Posterior Floor Plate (69% with SHH c1, 87.8% with SHH c5) and depleted hindbrain progenitors (Fig. 4D, top). Including FGF8 shifted the balance of regional fates back towards hindbrain progenitors (Fig. 4D, bottom), a finding that is consistent with the role of FGF8 at the Midbrain-Hindbrain Boundary (MHB) ^43^.

WNT inhibition and SHH activation also displayed interactive effects (Fig. 4E). XAV939 treatment (WNT inhibition) alone promoted diencephalic, neural crest and non-neuroectodermal fates in a dose-dependent manner, while SHH treatment alone induced progressive ventralization of anterior forebrain tissue, as previously described ^44^ (Fig. 4E, single morphogens). A combination of XAV939 and SHH showed synergistic effects leading to a higher proportion of non-neural ectodermal cells (Fig. 4E, morphogen combination). Notably, SHH was applied three days after XAV939, suggesting that SHH exposure might lead to the expansion of a pre-existing pool of non-neural ectodermal cells induced by WNT inhibition.

### hPSC lines and neural induction method impact patterning outcome

Different hPSC lines can have biases affecting differentiation capacity ^45^ and lead to neural organoids composed of different brain regions under unguided growth conditions ^7^. We tested if extrinsic patterning with morphogen pathway manipulation can override these biases by comparing cell type composition and gene expression patterns in control and morphogen treatment conditions from different PSC lines: WTC, HES3 and H9 (Fig. 5A-D and Fig. S9). We noticed strong differences in gross morphology between organoids from different cell lines generated using the same conditions (Fig. 5A). We summarized the gene expression profiles for each condition, cell line, and replicate experiment and found strong segregation of conditions and cell lines in a reduced dimensionality projection (Fig. 5B-C). Without morphogen treatment, we observed substantial variability in neural organoid regional cell types derived from different PSC lines (Fig. S9A-B). WTC iPS cells consistently generated organoids of telencephalic identity, H9 ES cells generated organoids with mixed telencephalic and retinal identities, whereas HES3 ES cells generated organoids with variable regional composition including telencephalic, diencephalic, retinal and neural crest identities (Fig. S9B). Under unguided conditions, none of the probed cell lines generated hypothalamic, hindbrain, mesenchymal, extraembryonic or endoderm cells (clusters 16-18; Fig. S9A-B).

**Fig. 5.**
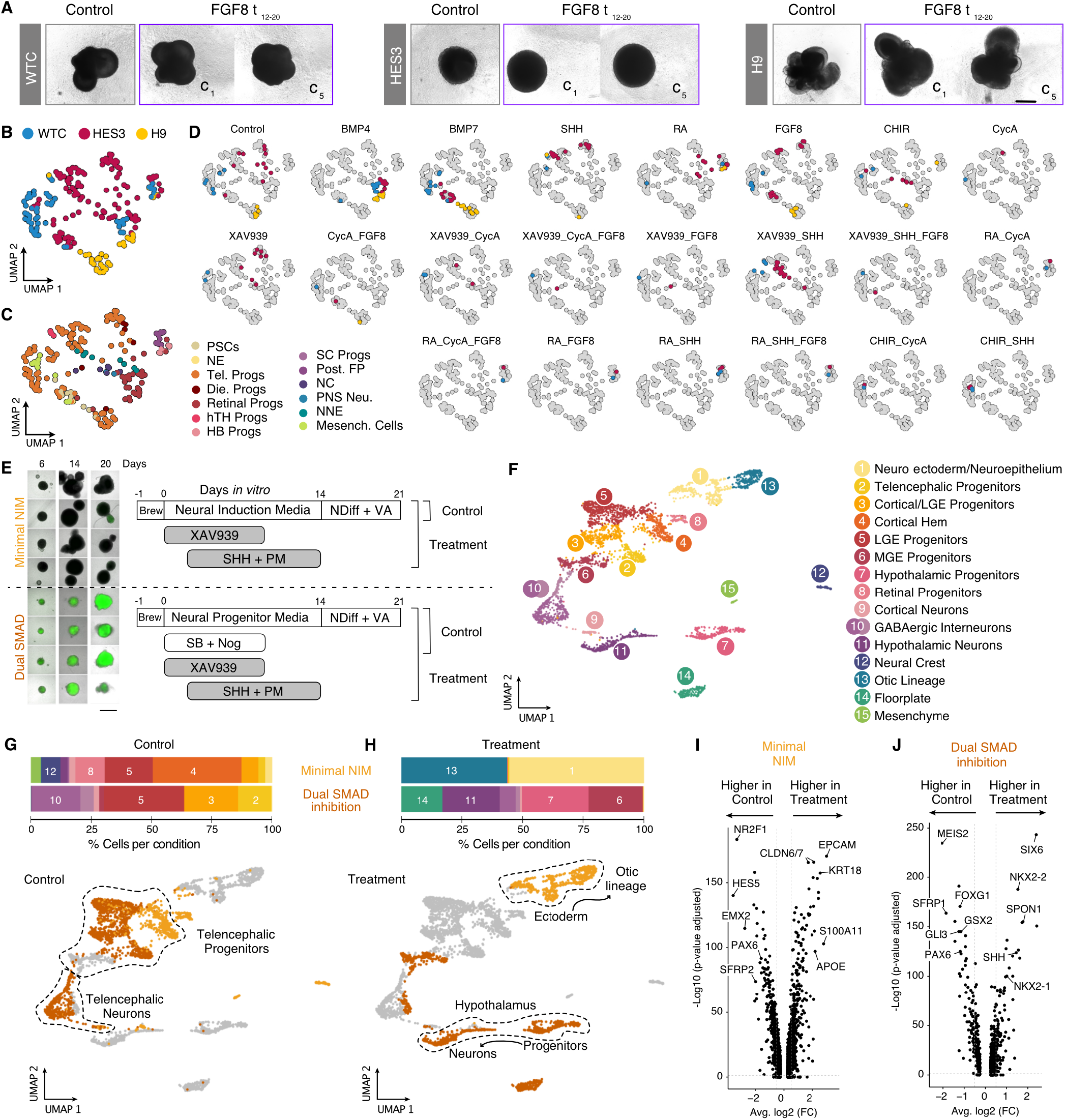
Morphogen patterning outcome is influenced by PSC line and neural induction media. (A) Brightfield images of day 20 neural organoids from WTC, HES3 and H9 lines in control conditions and under FGF8 treatment (days 12-20). Scale bar = 500 µm. (B,C) UMAP projection of summarized experiments, with each sample colored by cell line of origin (B) and predominant cluster identity (C). Each dot represents one pool of 4-16 organoids from the same line that were exposed to a certain morphogen treatment in each experiment/batch. Cluster identity legend: PSCs, Pluripotent Stem Cells; NE, Neuroectoderm; Tel. Progs, Telencephalic Progenitors; Die. Progs, Diencephalic Progenitors; hTH Progs, Hypothalamic Progenitors; HB Progs, Hindbrain Progenitors; SC Progs, Spinal Cord Progenitors; Post. FP, Posterior Floor Plate; NC, Neural Crest; PNS Neu, Peripheral Nervous System Neurons; NNE, Non-Neural Ectoderm; Mesench. Cells, Mesenchymal Cells. (D) UMAP projection split by morphogen identity (in some cases, including different treatment windows and concentrations) and colored by cell line of origin. (E) Organoid images on days 6, 14 and 20 of differentiation, cultured in the absence (top) and presence (bottom) of dual SMAD inhibition (SB431542 and Noggin). Treatment organoids were exposed to XAV939, Sonic hedgehog (SHH) and purmorphamine (PM) as indicated in the schematic (right). “Brew”, iPS Brew. Scale bar = 500 µm. (F) UMAP projection of the integrated samples (Control and Treatment from Minimal NIM and Dual SMAD organoids) colored, numbered, and labeled by cluster identity. (G,H) Organoid composition bar plots of control vs treatment conditions in each neural induction method (top) and UMAP projection with cells only from control or treatment conditions colored by neural induction method (bottom). (I,J) Differentially expressed genes between control and treatment conditions under minimal NIM (I) and dual SMAD inhibition (J).

Next, we assessed the differential response of PSC lines to different morphogen treatments (Fig. 5C-E). Although each morphogen pushed towards the generation of particular cell types, we observed cell line-specific responses for BMP4, BMP7, and FGF8 morphogen treatment conditions that seemed determined by their “default” identities (Fig. S9C and S9E). Focusing on exposure to SHH in the presence and absence of XAV939, we found that the three PSC lines generated hypothalamic progenitors at differential proportions. While WTC-derived organoids contained up to 80% hypothalamic progenitors, H9- and HES3-derived organoids were composed of less than 50% of this progenitor population (Fig. S9D). Hence, SHH-mediated ventralization showed differential efficiency in the three PSC lines. In summary, each cell line responded to morphogen treatments in the expected direction of the patterning cue, however, the efficiency or particular identity reached in this treatment window varied significantly between lines.

Neural organoid differentiation protocols apply different methods for the initial neural induction of the embryoid bodies including dual SMAD inhibition ^46^ as well as minimal neural induction media ^9^. We therefore designed an experiment to test if organoid ventralization using SHH and PM is impacted by different neural induction methods (Fig. 5E). We cultured organoids in minimal neural induction media or neural progenitor media (dual SMAD inhibition), supplied with XAV939, SHH and PM and performed single-cell RNA sequencing of each condition after 21 days of culture. We leveraged the hESC NKX2-1 reporter line (HES3, NKX2.1GFP/w) ^47^ as an indicator of ventralization, and observed differential activation of NKX2-1 in the two neural induction conditions, with strong NKX2-1 expression under dual SMAD inhibition but not in minimal neural induction media. Single-cell transcriptomic analysis showed a differential cell type abundance between the two conditions (Fig. 5F-H). While control organoids under both neural induction methods mainly contained telencephalic cell types (Fig. 5G), treatment with XAV939, SHH and PM resulted in different outcomes depending on the induction media (Fig. 5H). Dual SMAD inhibited organoids showed strong ventralization into floor plate and hypothalamus fates, whereas organoids in minimal neural induction media generated cell types of the otic lineage (Fig. 5G-H). Differential gene expression analysis supported this differential response (Fig. 5I-J). We suspect that deviation from the CNS lineage may be due to higher levels of BMP signalling in organoids grown in minimal neural induction media thereby promoting other ectodermal fates ^48^.

### Morphogen gradation influences dorso-ventral fore-brain proportion

Morphogen gradients form boundaries and organizing centers, which in turn influence region proportion during brain development ^3^. We used two approaches to test if gradual changes in morphogen exposure can control the proportion of dorsal and ventral telencephalon cells within complex human neural tissues (Fig. 6A-B). In a first approach, we use a two input microfluidic tree gradient generator, which has previously been used for antero-posterior neural tube specification (Microfluidic STem cell Regionalization, MiSTR) ^49^, to introduce a dorsal-ventral gradient. Here, a three-dimensional sheet of developing neural tissue received a gradient of SHH agonist and WNT antagonist molecules (Fig. 6B, top left). In a second approach, we exposed neural organoids to discrete concentrations of the same agonists representing different locations of the gradient, including a control condition not exposed to patterning molecules (Fig. 6B, top right). After 21 days in culture, we harvested MiSTR tissue and neural organoids to perform scRNA-seq, using cell hashing to track the MiSTR segment and organoid condition (Fig. 6B, middle to bottom). Datasets were merged and integrated using Harmony, RPCA, CCA, and CSS, with all integrations yielding a strong overlap between the two approaches (Fig. S10C). We note an exceptional FOXA2+ floor plate cluster, which was enriched in organoids treated with high doses of SHH (Fig. 6C-E and Fig. S10C).

**Fig. 6.**
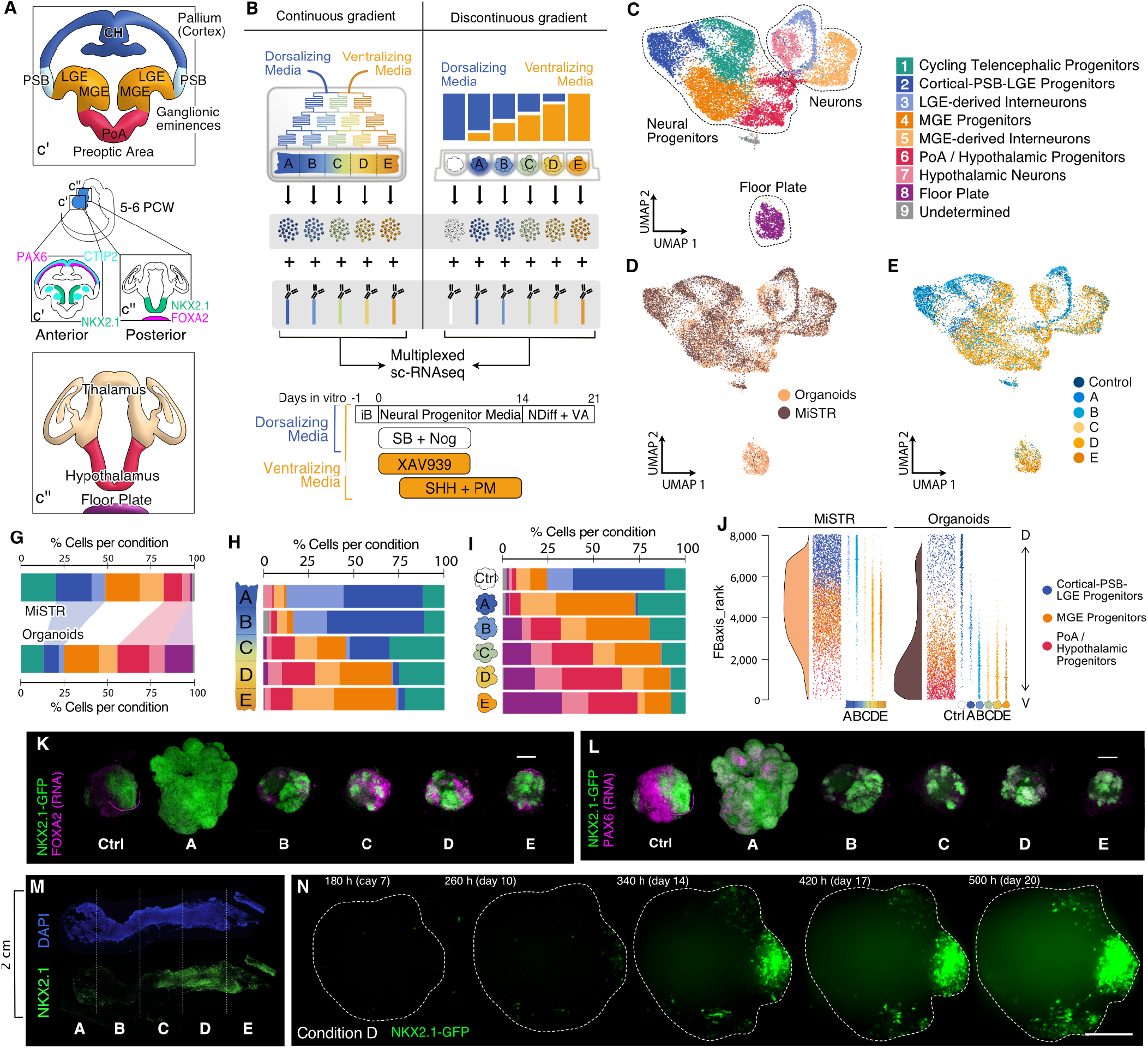
Morphogen gradients and discontinuous morphogen steps similarly recapitulate dorso-ventral patterning of the human forebrain. (A) Overview of dorsoventral patterning of the developing forebrain and the corresponding regions and marker genes expressed in each of them. Different antero-posterior sections and their patterning profiles are depicted in c’ and c”: telencephalic (FOXG1+, anterior) and hypothalamic (NKX2.1+, posterior) regions. (B) Simplified schematic of the microfluidic gradient tree (MiSTR) and the 96-well setup (organoids) used for treating developing neural tissue with increasing concentrations of SHH. Bottom, culture protocol for both experimental setups. (C-E) UMAP projection of Harmony-integrated single cell-RNAseq data from MiSTR and organoid experiments, colored by cluster (C), experimental setup (D) and section of origin (E). Annotations: PSB, Pallial-Subpallial Boundary; LGE, Lateral Ganglionic Eminence; MGE, Medial Ganglionic Eminence; PoA, Preoptic Area. (G) Cell type composition of all aggregated segments from each experimental setup. (H,I) Cell type composition of each segment from MiSTR tissues (H) and separately patterned organoids (I) (cell line = H9). (J) Forebrain axis score ranking of cells from each experimental setup, representing shifts from anterior-dorsal identities (higher ranks) towards posterior-ventral identities (lower ranks). For each of them: left, ridge plot showing distribution of cells along FBaxis_rank values; middle, jitter plot showing the distribution of each progenitor type along the FBaxis ranking (most dorsal cells at the top, most ventral cells at the bottom); right, same jitter plot showing separately each MiSTR segment and organoid condition. (K,L) HCR stainings of organoids patterned with increasing concentrations of SHH, PM and XAV939. Endogenous NKX2.1-GFP protein is shown in green, while FOXA2 (K) and PAX6 (L) mRNA levels are shown in magenta. (M) Immunohistochemistry staining of MiSTR neural tissue exposed to a continuous morphogen gradient. Orientation of the tissue is shown with A-E, indicating each subsection. (N) Still images from long-term light sheet imaging of organoids containing the NKX2.1:GFP reporter showing focal induction and spreading of ventral domains. Organoids were cultured using the concentration of SHH, PM and XAV9393 corresponding to condition D.

Similar cluster compositions were observed using both approaches, although with different proportions, and organoids contained more ventral regions than the tissue patterned with a microfluidic gradient (Fig. 6G-I). Low SHH regions in the MiSTR tissue produced cortical/lateral ganglionic eminence (Ctx/LGE) progenitors and neurons, whereas the lowest concentration of SHH in the organoids induced more ventral fates, including medial ganglionic eminence (MGE) progenitors and MGE-derived interneurons (FOXG1+, NKX2-1+). While increasing SHH concentrations induced mainly MGE fates in the MiSTR, high SHH induced a combination of hypothalamic fates (FOXG1-, NKX2-1+) and diencephalic floor plate fates (FOXA2+) in the organoids.

Forebrain axial scoring and ranking (FBaxis_rank) revealed a continuity of the axial domains generated in each of the settings, with MiSTR tissue cells dominated by telencephalic dorso-ventral fates and organoid cells dominated by diencephalic ventral fates (Fig. 6J and Fig. S10D). Immunostaining and HCR in situ hybridization confirmed these findings, revealing NKX2-1 expression in MiSTR segments with higher SHH exposure (Fig. 6M, sections C-E), whereas organoids displayed NKX2-1 expression in all conditions, including Control (Fig. 6K-L). Overall, this data indicates a higher proportion of ventral cells in the SHH-stimulated organoids compared to the MiSTR, as well as a slightly more caudal identity in the organoid setting. We speculate that these differences could be due to differences in morphogen availability in a flowing microfluidic PDMS chamber versus a plate setting with intermittent media change. Interestingly, live fluorescent imaging of NKX2-1:GFP organoids growing in high SHH conditions revealed the localized emergence of NKX2-1 driven fluorescence ^50^ despite non-localized exposure to SHH, and these foci gradually spread (Fig. 6N). Meanwhile, rectangular monolayer cultures showed homogeneous, low expression levels of NKX2.1-GFP on day 5 that progressively increased in a simultaneous manner across the tissue (Fig. S10E). Altogether, these data show that graded exposure to morphogens can be harnessed to influence brain region proportion in complex 3D neuroepithelial tissues, and points to interesting differences in self-organizing patterning dynamics in globular versus planar neuroepithelial tissues.

## Discussion

Stem cell-derived neuroepithelium generated in vitro presents a unique opportunity to systematically study how modulation of morphogen signaling pathways impact a naive neuroepithelium. In this study, we provide a thorough evaluation of timing- and concentration-dependent effects of morphogen signaling modulators on developing human neural organoids. We delimit competence windows for SHH, FGF8, RA, BMP4/7 and WNT pathway regulation to control the induction of different types of neural and non-neural fates. We also demonstrate that cell fate decisions can be controlled by modulating morphogen concentrations, as illustrated by SHH and RA, where we observe enrichment of different regional fates at low and high doses in comparison to untreated organoids. In particular, RA showed the strongest effects on neural organoid patterning, both across replicates and cell lines, generating a wide variety of regions at different time points and concentrations. In addition, we dissect synergies of morphogens that are known to cooperatively pattern the developing neural tube. For example, RA stimulated the production of either posterior floor plate in combination with SHH or hindbrain progenitors in the presence of FGF8. The interaction between FGF8 and RA is consistent with a known role in the specification of hindbrain compartments ^2,3^, while the synergy between SHH and RA has only been reported in spinal cord ^51–55^ and other tissues ^56,57^. Another example of a novel interaction involves WNT inhibition (XAV939) and subsequent SHH activation, which together led to an increase in non-neural ectoderm. This effect could only be observed in the absence of dual SMAD inhibition, highlighting the relevance of neural induction methods for morphogen patterning outcomes.

We also observed divergent responses to morphogen exposure resulting from different hPSC lines. This lack of consistency is likely due to genetic or epigenetic differences between cell lines that can lead to different baseline expression of or susceptibility to WNT and other developmental signaling pathways ^45^. Recent work has evaluated somatic mutation effects on neuronal differentiation competency from different iPSC lines and provided genotyping resources for hundreds of hESC lines ^58,59^. These resources, together with high-throughput and systematic screens will help understand the nature of differences between hPSC lines to generate human neural tissues.

An important finding in our study is that most tissues assayed under a great diversity of conditions were composites of multiple regional identities. Interestingly, regions that were prevalent in each condition are known to be spatially adjacent in the primary developing human brain. This suggests that each organoid has a self-organizing response to a static condition such that cells differentially interpret cues resulting in relevant regional boundaries. There has been substantial progress in developing protocols that generate specific brain regions from human pluripotent stem cells, yet in the majority of protocols there is incomplete patterning to a singular region of the brain ^60^. Further analyses are needed to understand the cellular and tissue mechanisms underlying regional boundary formation. It is likely that gradients emerge in the organoids to establish these regional domains. Indeed, directly controlling morphogen exposure using gradient microfluidic experiments showed that there is a shifting of composition proportion along the concentration axis.

In addition, the neural induction method has a substantial impact on the resulting organoid regionalization. Strong dual SMAD inhibition-based neural induction has the advantage of producing predominantly central nervous system fates, whereas gentler approaches can produce tissues of multiple neuro- and non-neuroectodermal origins, which could be harnessed to engineer interactions between the neural crest, placodes, and parts of the brain.

Altogether, our work provides an integrated overview of morphogen-induced regionalization of neural tissues in vitro and enhances our understanding of morphogen-driven patterning in the early human embryo. It also demonstrates, together with recent work on later stages of neural organoids ^61^, that morphogen screens with single-cell genomic readouts provide exciting opportunities to develop and optimize new protocols through systematic exploration of organoid patterning.

## Supporting information

Sup_cluster_markers_1

Sup_cluster_markers_2

Sup_cluster_markers_3

Sup_regulons_BMP4

Sup_regulons_BMP7

Sup_regulons_SHH

Sup_regulons_FGF8

Sup_regulons_RA

Sup_regulons_CHIR

Sup_regulons_XAV939

## AUTHOR CONTRIBUTIONS

F.S.C. designed and performed the experiments, analyzed and interpreted the results, and wrote the manuscript. A.J. executed and analyzed long term light-sheet imaging experiments and contributed to HCR establishment and experimental design. Z.H. performed and supervised computational analyses. R.O. performed hPSC, organoid culture and cell hashing experiments. C.R., P.R. and G.S.R. performed MiSTR experiments. M. Santel performed organoid culture and cell hashing experiments. J. J. performed SCENIC analysis and J.F. performed organoid cell type projection to the developing human brain reference. M. Seimiya performed HCR stainings. A. K. designed and supervised MiSTR experiments and provided feedback on the entire project. J.G.C. and B. T. supervised the project, designed the experiments, interpreted the results and wrote the manuscript.

## Declaration of interests

The authors declare no competing interests.

## ACKNOWLEDGEMENTS

We thank the Treutlein, Camp and Kirkeby labs for insightful discussions and feedback. We also thank Eliane Duperrex for helping to establish in situ HCR for this project. We thank the Genomics Facility Basel and Single Cell Facility for technical support in scRNA-seq and imaging experiments. We also want to thank Prof. Ed Stanley and Prof. Andrew G. Elefanty (MCRI and Monash University) and the Kirkeby lab for kindly sharing the HES3, NKX2.1GFP/w cell line. We thank Sophie Seidel for fruitful discussions on regulon-based analysis, and Patricia L. Murphy for help with manuscript formatting. Work in the laboratory of B.T. is supported by the European Research Council (organomics and braintime (to B.T.)); the Chan Zuckerberg Initiative DAF, an advised fund of the Silicon Valley Community Foundation (grant no. CZF2019-002440); the Swiss National Science Foundation (grant no. 310030_192604); and the National Centre of Competence in Research Molecular Systems Engineering. J.G.C. was supported by the European Research Council (Anthropoid-803441) and the Swiss National Science Foundation (project grant 310030_84795) and A.K. by the Novo Nordisk Foundation (NNF18OC0030286 and NNF21CC0073729).

## Materials and Methods

### iPSC and ESC culture

NKX2.1GFP/w hESCs were obtained from Agnete Kirkeby’s research group at the University of Copenhagen, after the arrangement of an MTA with Prof. Ed Stanley and Prof. Andrew G. Elefanty (Murdoch Childrens Research Institute, Melbourne). H9 (WA09) hESCs were obtained from WiCell and WTC hiPSCs from the Allen Institute for Cell Science. Stem cells were grown at 37°C, 5% CO2 in feeder-free conditions, on 6-well tissue culture plates coated with hESC-Qualified Matrigel (Corning). They were fed every other day with mTesR Plus (StemCell Technologies) or StemMACS iPS Brew XF (Miltenyi) and passaged when reaching around 80% confluence with EDTA for gentle dissociation. All lines were tested regularly for copy number changes using the Agilent ISCA 8×60K v2 array and no abnormalities were detected. Additionally, a karyotyping control was performed and showed normal XX (NKX2.1GFP/w and H9) and XY (WTC) karyotypes. These analyses were performed by the Cell Guidance Systems Genetics Service (Cambridge, UK). Regular PCR-based Mycoplasma testing (Biological Industries) was performed to discard potential Mycoplasma infections.

### Neural organoid generation and incubation with patterning molecules

Stem cells were washed with PBS once and trypsinized using TrypLE™ Express Enzyme (Thermo Fisher Scientific) to obtain single-cell suspensions and 500 cells were plated in each well of ultra-low attachment 96-well plates (day 0). A 1-5 min centrifugation of the well-plate was performed at 200g and cells in suspension were cultured for one day in mTesR Plus (StemCell Technologies) to allow for aggregation and Embryoid Body (EB) formation. On day 1, EBs were transitioned to Neural Induction Media (NIM, containing DMEM/F12, 1% N2 supplement (v/v), 1% GlutaMAX supplement (v/v), 1% MEM-NEAA (v/v) and 1 µg/mL Heparin). Neural Induction Media was maintained for 14 days and then switched for NDiff+VA, with the exception of the RA/XAV939/SHH/FGF8 timing experiments, which used NDiff+VA instead until day 21. The specific morphogen treatment windows and concentrations used per condition are specified in Table S1 and S2.

In the minimal NIM vs Dual SMAD inhibition experiments, control Dual-SMAD organoids were cultured in Neural Patterning Media (see MiSTR and MiSTR-like organoid culture sections below) and exposed to 10 µM SB (SB432542, Miltenyi) and 100 ng/mL Noggin (rh-Noggin, Miltenyi) from day 0 to 9. Minimal NIM organoids were cultured in the Neural Induction Media (NIM) described above. For organoid patterning, we used 4.5 µM XAV939 (Miltenyi) from day 0 to 9 and 180 ng/mL SHH (rh-SHH/C24II, Miltenyi) together with 270 nM Purmorphamine (Miltenyi) from day 3 to 14.

**Table S1.**
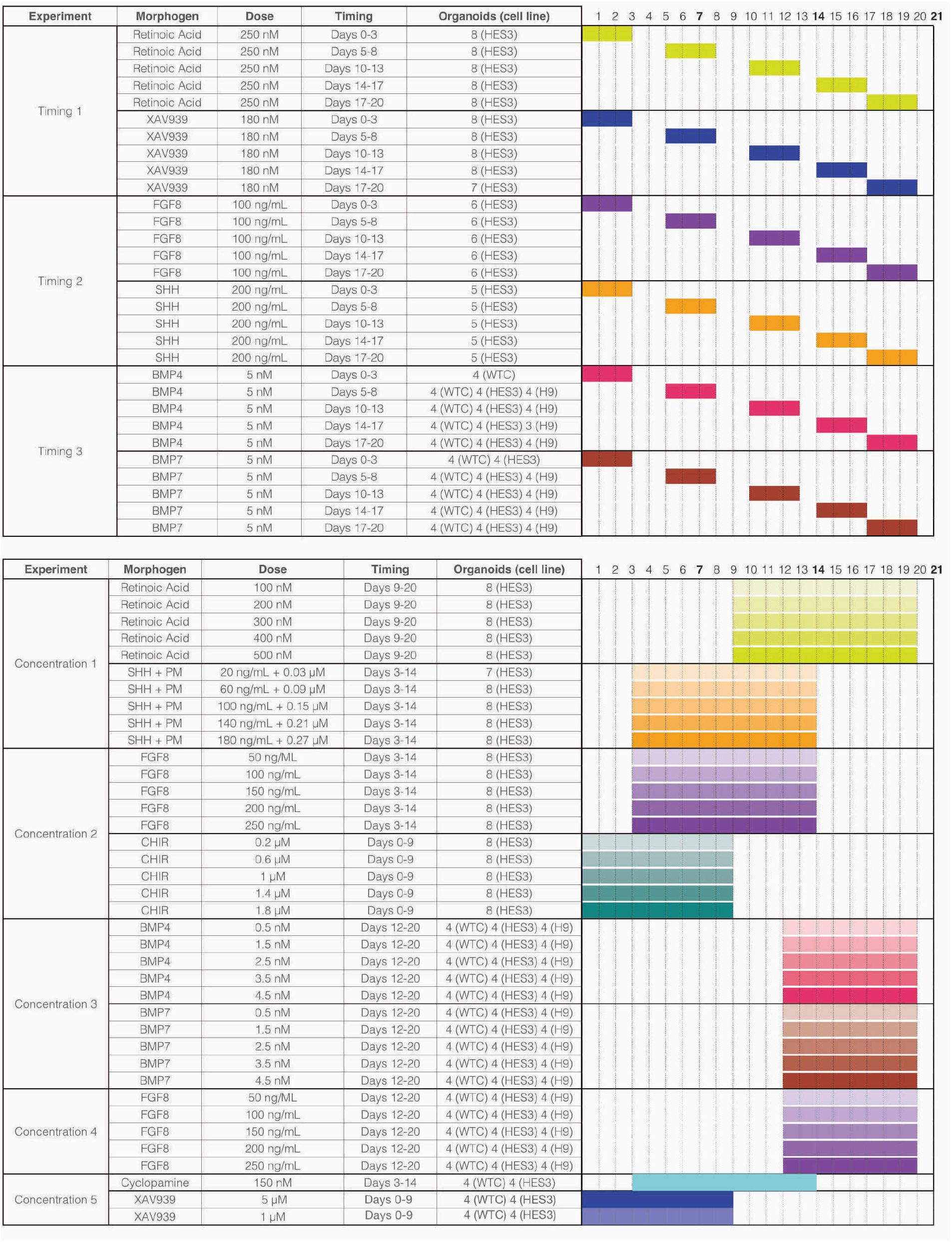
Experimental conditions for the probing of morphogen timing and concentrations in human neural organoids. The number of assayed organoids per morphogen treatment from each ESC/iPSC line is indicated in the last column. The right section of the table depicts a summary of the patterning protocol for each condition, with the culture days indicated at the top. Color intensity represents morphogen concentrations (higher intensity means higher concentration).

**Table S2.**
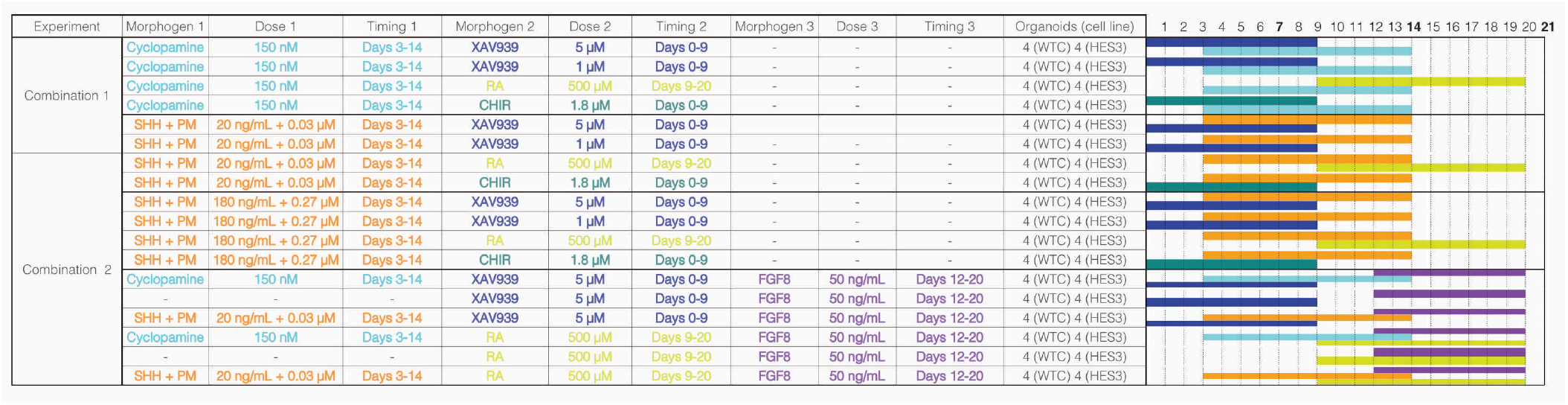
Experimental conditions for the probing of morphogen combinations in human neural organoids. Up to three morphogen pathways were targeted in the same treatment; their dose and timing is stated in separate columns. The number of assayed organoids per morphogen treatment from each ESC/iPSC line is indicated in the last column. The right section of the table depicts a summary of the patterning protocol for each condition, with the culture days indicated at the top.

### MiSTR cultures

MiSTR culture was performed as previously published ^49^ with some modifications to pattern in the dorsoventral axis. One day before the assembly of the MiSTR device and start of differentiation (day -1), H9 cells were detached from the well using EDTA to obtain single-cell suspensions, which were adjusted to a concentration of 1.25 × 10^6^ cells/mL in iPS Brew + 10 µM ROCK inhibitor (Y-27632, VWR). 800 µL of cell suspension (approximately 1 million cells) were plated on top of a pure GFR-Matrigel (Corning) bed polymerized within a PDMS chamber in the bottom part of the MiSTR device. Cells were incubated for a day (37ºC, 5% CO2) to allow them to proliferate and form a uniform sheet. On the next day, dead cells and debris were removed from the chamber by washing twice with 1:20 KOSR (Knock-Out Serum Replacement, LifeTech) in DMEM/F12 (LifeTech). After aspirating bubbles and pre-perfusing the microfluidic tree with the same solution, the top part of the MiSTR device (containing the PDMS microfluidic tree) was fit over the bed of ESCs. Lastly, the polycarbonate lid was placed on top and the whole setup was pushed into the POM cassette for stabilization.

Once the “sandwich” was assembled, two high-precision GasTight syringes (Hamilton) with different media were attached to the tubing coming out of either side of the MiSTR device:

- The left side syringe contained default Neural Patterning Media: 50% DMEM/F12 (LifeTech), 50% NeuroMedium (Miltenyi), 1:200 Glutamax (LifeTech), 1:250 P/S (Penicillin/Streptomycin, LifeTech), NB-21 (NeuroBrew-21 without vitamin A, Miltenyi) and 1:200 N2 supplement (LifeTech). This media was complemented with SB (SB432542, Miltenyi) and Noggin (rh-Noggin, Miltenyi) for neural induction from day 0 to day 9.
- The right side syringe contained the same Neural Patterning Media as above, also complemented with SB and Noggin for neural induction on days 0-9, and the following patterning molecules: from day 0 to 9, 5 µM of XAV939 (Miltenyi); from day 3 to 14, 200 ng/mL of SHH (rh-SHH/C24II, Miltenyi) and 0.3 µM of PM (Purmorphamine, Miltenyi).

After bringing the MiSTR device into the incubator and assembling the syringes on high-precision neMESYS pumps, the flow was initiated and kept at a constant rate of 160 µL/h. Meanwhile, the outlet tubing would collect waste media into a small glass bottle placed on an upper shelf of the same incubator. The syringe media was changed or replenished every 2 or 3 days and the differentiation was maintained in the microfluidic tree for 14 days. Then the MiSTR device was disassembled, and the neural sheet was recovered and completely embedded in GFR-Matrigel by adding 200 µL on top of it. After polymerization of the fresh Matrigel, the tissue was transported into a Petri dish with Ndiff+VA media (see composition above, under “Neural organoid generation and incubation with patterning molecules”). Cultures were kept in the incubator under mild rocking conditions to ensure good perfusion and media was changed every 3-4 days until day 21. On day 21 of MiSTR culture, the neural sheet was retrieved and cut in 5 equal parts from left to right, following the longitudinal axis of the gradient in the chamber. A custom-made mould was used to standardize the sizes of each piece. From left to right (basic NPM on the left to NPM with patterning factors on the right), the sections were called A, B, C, D and E. Each section from A to E is represented by 2 independent MiSTR experiments, which were pooled together for dissociation. Before and after each experiment, the different parts of the MiSTR device were sterilized appropriately by either ethanol baths or autoclaving, according to their material.

### MiSTR-like organoid cultures

On day -1, EB aggregation of H9 and HES3 (NKX2.1GFP/w) hESCs was performed as described in the previous section, using iPS Brew and Rock inhibitor at 1:200. For neural induction, Neural Patterning Media was employed together with SB and Noggin-mediated dual SMAD inhibition from day 0 to day 14. After this, media was changed to Neural Differentiation Media with vitamin A (composition previously described) until day 21. Overall, organoids were grown in identical media conditions as the MiSTR cultures, but in 96-well plates for the entire time course. Each of the organoid conditions was treated with the average morphogen concentration to which the corresponding MiSTR segment was exposed, as shown in Table S3.

**Table S3.** Culture conditions for MiSTR-like organoids. The morphogen dose given to each condition is stated in columns Control-E.

| Timing | Morphogen | Control | A | B | C | D | E |
| --- | --- | --- | --- | --- | --- | --- | --- |
| Day 3-14 | SHH (ng/mL) | 0 | 20 | 60 | 100 | 140 | 180 |
| Day 3-14 | PM (nM) | 0 | 30 | 90 | 150 | 210 | 270 |
| Day 0-9 | XAV939 (μM) | 0 | 0.5 | 1.5 | 2.5 | 3.5 | 4.5 |

### Live imaging of developing MiSTR-like organoids and 2D neural sheets

For the light-sheet imaging experiments, MiSTR-like organoids from condition “Control” (n=4) and “D” (n=4) were aggregated on day -1 and transferred to an imaging cuvette one day later (Day 0 of the culture protocol). Matrigel diluted to 1:50 was used to fix the organoids in their position in the microchambers within the cuvette. The LS1 Live light-sheet microscope (Viventis) was used to image each organoid every hour with a 25x objective demagnified to 18.5x, across 200 optical slices separated by 2 µm step size. In total, MiSTR-like organoids were imaged for 21 days ^50^.

For the imaging of 2D cultures of MiSTR-like neural progenitors, a custom-made chamber was made with the same dimensions as the MiSTR chamber. This chamber was used to contain 1 million H9 and HES3 (NKX2.1GFP/w) hESCs, which were seeded on day -1 on a bed of pure Matrigel. From day 0 until day 14, they were exposed to NPM and the morphogen cocktail corresponding to “Control” and condition “D” (as described in the previous section). Imaging of the resulting neural sheets was performed using a Nikon Ti2 microscope with a spinning disk module. Tile scans of the entire tissue were taken every day until NKX2.1-GFP activation, at 4x magnification in 3 different z-step locations. Media was exchanged every day and the cultures were maintained until day 21.

### Organoid dissociation

On day 21 of organoid culture, all the organoids from the same conditions were collected from their individual well and pooled for dissociation together. A-E segments of 21-day-old MiSTR tissue were processed in the same way as organoids.

The Miltenyi neural tissue dissociation kit was consistently used in all experiments. First, tissue from each dissociation pool (further referred to as “sample”) was transferred to a 5 mL Eppendorf tube, where it would be washed twice with 1 mL of HBSS without Ca2+ and Mg2+ (HBSS w/o, Thermo Scientific). After removing the supernatant, 1 mL of enzyme mix 1 (Enzyme P/Papain + Buffer X from Miltenyi Neural dissociation kit) was added to each sample and incubated at 37ºC in the water bath or inside the incubator for 15 min. In the case of the MiSTR tissue dissociations, the gentleMACS Octo Dissociator (Miltenyi) was used inside the incubator for a milder and more progressive homogenization of the tissue into single cells.

After the first incubation step, 15 µL of enzyme mix 2 (Enzyme A/DNAse + Buffer Y from Miltenyi Neural Dissociation kit) were added and the sample was mixed carefully with a wide bore pipette, then triturated 5-10x with a p1000 pipette and triturated again 5-10x with a p200 before another 15 min of incubation at 37ºC. Samples were then further triturated with a p200 and incubated for another 10 min at 37ºC. If there were still clumps, samples were triturated once more with a p200. At this stage, the quality of the dissociation was monitored by observing 1 µL of cell suspension under the microscope at 10X. If there were still clumps, further 5 min incubation and trituration steps would be performed. If the suspension was homogeneous, we would proceed to a filtering step.

For cell suspension filtrations, 20 µm PluriSelect cell strainers were used. These were placed on top of a new 5 mL tube and pre-wet with 500 mL of HBSS w/o before passing the cell suspension through it and washing any trapped cells with 1 mL of HBSS w/o and 1 mL of 0.5% BSA (Miltenyi). From these filtered cell suspensions, 20 µL were taken from each of them to count cells and check their viability. In the meantime, samples were centrifuged once for 5 min at 300g using a swing centrifuge to minimize cell loss by concentrating the pellet at the bottom of the tube. After this step, the supernatant was removed, and cells were resuspended in staining buffer (0.5% BSA in PBS-/-) at a concentration of 7,777 cells/µL.

### Cell hashing (CITE-seq)

Once all samples were ready at the desired concentration, 45 µL of cell suspension was transferred to a new 1.5 mL Eppendorf tube (approximately 350,000 cells) and 5 µL of Human TruStain FcX™ Fc Blocking reagent (Biolegend) were added to block unspecific binding of the hashing antibodies. After mixing, the suspension was incubated for 10 min at 4ºC. Right after blocking, 2 µL (1 µg) of the corresponding hashing antibody (Totalseq-A, Biolegend, Table S4) were added to the blocked samples, mixed with a pipette, and left to incubate for 30 minutes at 4ºC. Every 10 minutes the tubes were mildly shaken to avoid precipitation of cells at the bottom of the tube. When the incubation time was finished, 1 mL of staining buffer was added to dilute each sample and tubes were spun down in a swing centrifuge for 5 mins at 300g and 4ºC. The supernatant was then removed, and an extra washing step was performed, during which 20 µL of cell suspension were taken out for cell counting and viability checking. After the last washing step, cell pellets were resuspended in staining buffer at a concentration of 1,000 cells/µL. Once all samples were adjusted in concentration, 6-12 samples were pooled together 1:1, usually by mixing 10 µL of each of them. The suspension was kept on ice during 10X preparations and 20-25 µL of the hashed pool (20-25,000 cells) were loaded in one lane of the Chromium chip (10X Genomics).

**Table S4.**
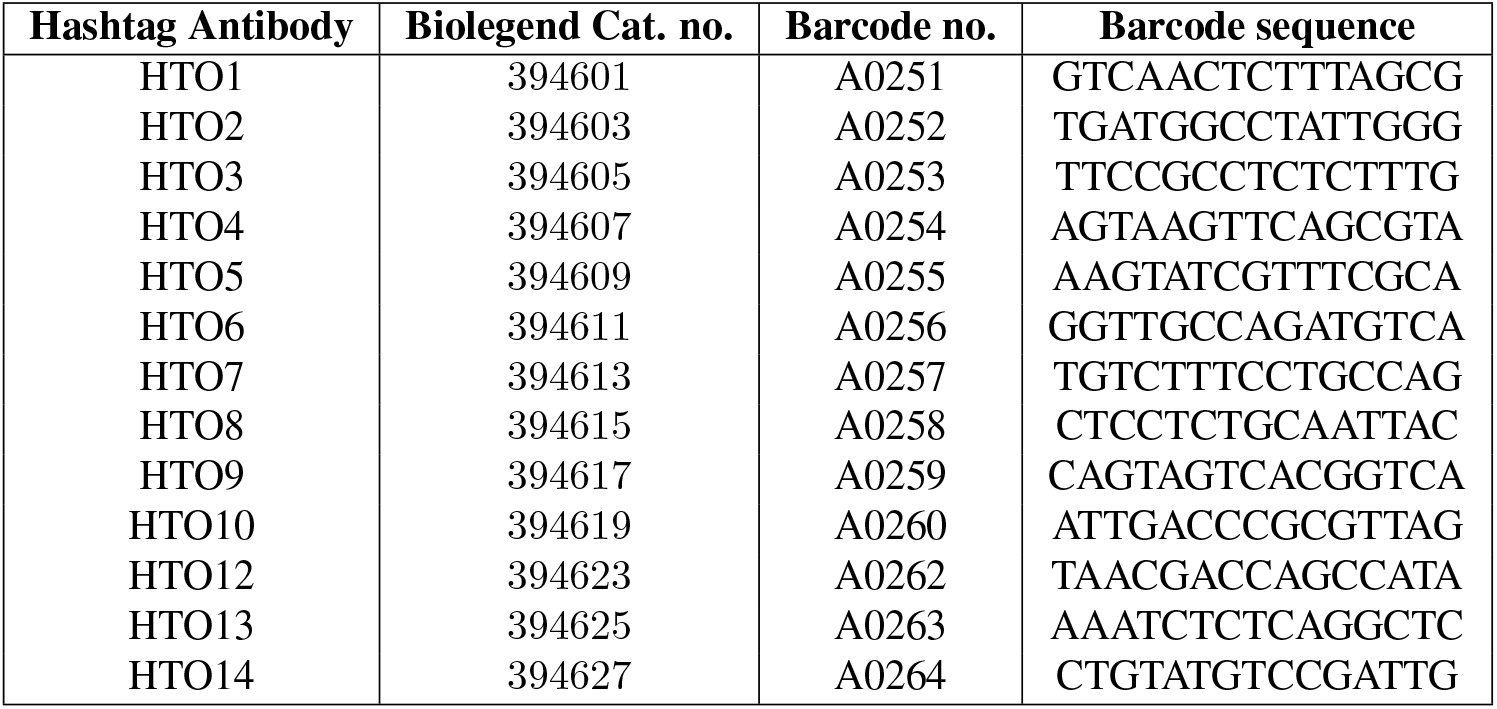
List of all the Totalseq-A antibodies used for cell hashing and their corresponding barcode sequence.

### Single cell transcriptome (cDNA) library generation

We used the Chromium Next GEM Single Cell 3’ v3/3.1 kit (10X Genomics) to generate single cell transcriptome libraries from hashed samples. The procedure followed the steps listed in the corresponding 10X protocol (CG000183 Rev A for the v3, CG000204 Rev D for the v3.1).

Briefly, the Chromium chip and Controller were used to encapsulate single cells in GEMs (nanoliter-scale Gel beads in EMulsion) together with a mastermix for reverse transcription (RT) of its messenger RNAs and a gel bead functionalized with polydT sequences that capture the cell’s mRNA. In addition, these gel beads also contain a UMI (Unique Molecular Identifier) that individually tags all original RNA molecules and a 10x barcode that labels all cDNAs belonging to the same cell. A Truseq read 1 adaptor sequence is also included to facilitate transcriptome sequencing in Illumina platforms. Once cells were encapsulated in GEMs, an RT step was run to convert polyadenylated mRNAs into full-length cDNA sequences with a UMI and cellular barcode. cDNA products were then purified with magnetic beads (Dynabeads MyOne SILANE, manufactured by Thermo Fisher but included in 10X Genomics kits) and further amplified in an extra PCR step using a partial TruSeq read 1 and partial TSO (Template-Switch Oligo) as primers. Another purification step based on construct size was performed with SPRIselect reagent (Beckman Coulter) prior to quantification and quality control with a Bioanalyzer High Sensitivity chip (2100 Bioanalyzer, Agilent).

One quarter of the cDNA from each 10X sample was carried over for gene expression library construction. In short, cDNA was fragmented to eliminate the TSO incorporated in the previous step and ends were repaired and A-tailed. Reaction products were purified with SPRIselect reagent and Truseq read 2 adaptors were ligated, one of them being a partial read to provide an overhang for the following PCR. Another cleanup step with SPRIselect was performed before the Sample Index PCR when P5 and P7 sequences (for Illumina sequencing) and sample indices (to allow for multiplexing in the sequencer) were included in the construct. The final product was purified with SPRIselect magnetic beads and a concentration and quality check were performed by running a Bioanalyzer High Sensitivity chip before sequencing.

### Single-cell hashtag (HTO) library generation

The first steps of HTO library preparation were done synchronously with the cDNA library until the cDNA amplification step of the 10x library workflow. From that point onwards, a modified version of the Biolegend protocol was used. The barcoded antibodies are coupled to oligo sequences called hashtags (HTOs), which contain a polyadenylated sequence at their 3’ end that enables hybridisation with the poly(dT) capture sequences in GEM beads. In parallel to endogenous mRNAs, HTO sequences were reverse-transcribed, and the resulting cDNA was amplified using as PCR oligos the partial TruSeq read 1 adaptor (provided in the 10x kit) and an additive primer (a partial TruSeq read 2, sequence provided by Biolegend and synthesized by Integrated DNA Technologies). The additive primer annealed on the 3’ end of the construct and enabled its exponential amplification.

During the post-cDNA amplification clean-up step, hashtag-derived fragments (HTO) were separated from mRNA-derived fragments (cDNA) based on their size (<180 bp) and processed separately to construct a hashtag-specific library. The HTO fraction was subjected to a Sample Index PCR where the P5 adaptor was added through the SI-PCR primer mix (10x kit) together with the P7 adaptor and an index sequence (both contained in a Truseq D70x_s oligo, sequence synthesized by Integrated DNA Technologies) to allow for simultaneous sequencing with other libraries. For each sample, reactions contained 5 µL of HTO fraction, 2.5 µL of SI-PCR primer (10 µM stock), 2.5 µL of TruSeq D70x_s primer (10 µM stock), 50 µL of Kapa Hifi Hotstart Ready Mix (Roche) and 40 µL of Nuclease-free water (Invitrogen). After the Sample Index PCR, a final SPRIselect-based cleanup was performed before resuspension in Nuclease-free water and library QC and quantification.

### Sequencing and genomic data preprocessing

The pooled libraries were appropriately diluted and sequenced at the Genomics Facility Basel (GFB). Sequencing data was demultiplexed by the GFB, using bcl2fastq version 2.20.422 with the following parameters: –ignore-missing-bcls –ignore-missing-controls –ignore-missing-positions –ignore-missing-filter –no-bgzf-compression –barcode-mismatches 1. 10x libraries were sequenced in SP/S1/S2 flowcells with Illumina NovaSeq technology, using paired-end 28/10/10/90 or 28/8/0/91 configuration (Read1/IDX i7/IDX i5/Read2). After sample demultiplexing, cell ranger v3.1.0 was used with GRCh38-3.0.0 as a reference to derive gene-cell count matrices from the sequencing read (fastq) files of the gene expression library. To obtain hashtag-cell count matrices, CITE-seq-Count v1.4.3^24^ was run on the hashtag library sequencing read files with the following parameters: -cbf 1, -cbl 16, -umif 17, -umil 28, and a variable number of targeted cells depending on the loaded cells per experiment (ranging from 12,000 to 16,000 cells).

### Demultiplexing of hashing antibody labels

The hashtag-cell count matrices obtained from CITE-seq-Count were preprocessed with a custom-made script, which involved transposition and column/row name adaptation. Once adapted, they were subset for the cell barcodes present in both hashtag and gene expression library and added as an “HTO” assay to normalized Seurat objects. They were then normalized using the NormalizeData function using the centered log-ratio transformation (CLR) method. After normalization, HTODemux was run on the “HTO” assay with a positive.quantile of 0.99-0.999 (adapted depending on the sample) to obtain a hashtag labelling classification of each cell. The quality of this classification was assessed by visualizing hashtag assignments on tSNE and RidgePlots.

### Single-cell gene expression data analysis

Gene count matrices from each experiment were pre-processed and analyzed using Seurat v4.3.0^62^. A quality control step was consistently applied by filtering out cells with gene counts lower than 1,000 and mitochondrial gene contents of 10% or higher. The count matrices were then normalized and scaled, either using the functions NormalizeData (logarithmic normalization) and ScaleData in the MiSTR experiments or SCTransform in the patterning screenings. In all cases, cell cycle scoring was performed to remove the effects of cell cycle variables in the data during the scaling step. In addition, a custom-made function was applied to ignore mitochondrial and ribosomal genes in downstream analysis.

Variable feature selection was performed with the “mean.var.plot” method or within SCTransform, and the obtained variable genes were used as input for PCA calculation. The PCs with correlation levels to cell cycle scores higher than 0.3 were excluded from downstream analysis. A variable number of the other PCs – determined by ElbowPlot visualization of their contribution to dataset variance - were used for further dimensionality reduction with t-SNE or UMAP embeddings and for neighbor finding and clustering. The clustering resolution and UMAP parameters were adapted for each dataset to retrieve the most meaningful biological information. In some intermediate steps, a list of relevant patterning genes was provided as input to PC calculation to enhance visualization of differences in patterning. Feature, dotplot, heatmap and violin plotting was performed with the package ggplot2 v3.4.1^63^, Seurat ^62^ and SCpubr v1.1.2^64^. Differential gene expression testing was performed with Seurat’s function FindMarkers and the wilcoxauc function from the presto package ^65^.

Once each experiment was analyzed separately, the single-cell gene/hashtag data was merged with Seurat or integrated using Harmony ^66^, CCA/RPCA ^67^, or CSS/RSS ^25^. In RSS integration, a pseudo-cell aggregated version was used as a reference. The LabelTransfer function from Seurat was used to project the gene expression data to the developing mouse ^68^ and human brain ^26^ single-cell transcriptome datasets and the VoxHunt package (v.1.0.1) ^69^ was used for spatial projections to the developing mouse brain, using the Allen Brain Atlas ISH reference.

### In situ Hybridization Chain Reaction (HCR)

Tissue collection and fixation: on day 21 of culture, neural organoids were collected from the 96-well plates and pooled by condition in 2 mL Eppendorf tubes. After removing leftover media, 2 mL of cold 4% PFA (Thermo Scientific, 28908) in RNAse-free PBS (Invitrogen, AM9625) was added and samples were fixed for 2 hours or overnight at 4ºC. All subsequent incubation steps were done with mild rocking using a nutator at 4ºC unless otherwise stated to preserve the quality of endogenous mRNAs. After fixation, PFA was removed and samples were washed 3 times with PBST (RNAse-free PBS and Tween20, Sigma Aldrich) with incubation times of 15 minutes. After the last wash, half of the PBST was removed and the same amount of methanol (MeOH) was added to make it a 50/50 PBST/MeOH mixture (v/v). Samples were incubated for 10-15 minutes and the PBST/MeOH solution was exchanged for 100% MeOH, then incubated for 15 minutes and replaced by new, pure MeOH for overnight storage at -20ºC.

On the day of the experiment, samples were rehydrated with a series of graded PBST/MeOH washes (25% PBST / 75% MeOH, 50% PBST / 50% MeOH, 75% PBST / 25% MeOH, 100% PBST twice) for 5 minutes each at 4ºC. Then PBST was removed, and samples were treated with 0.5 mL of 10 µg/mL proteinase K (AM2546, Thermo Fisher) in PBST for 3 minutes at room temperature. This concentration and time were previously optimized for good permeabilization without disruption of neural organoid tissue. Next, organoids were washed twice for 2 minutes with 1.5 mL of PBST, then postfixed with 0.5 mL of 4% PFA (20 min at RT) and washed again 3 times for 5 minutes with 1.5 mL of PBST. Detection stage: Before hybridization, organoids were treated with 200 µL of probe hybridization buffer for 30 min at 37 ºC inside a Thermomixer covered by the ThermoTop (Eppendorf) to avoid evaporation of the buffer. The probe hybridization buffer was then removed and 200 µL of probe solution were added to the organoids, which were incubated overnight at 37ºC as previously mentioned. The probe solution contained 100 µL of probe hybridization buffer and 0.5 µL of each hybridization probe (targeting each gene, 1 µM stock), up to a total of 5 probes. On the next day, excess probes were removed by washing 4 times for 15 min with pre-heated probe wash buffer (30% formamide, 5x SSC, 9 mM citric acid pH 6, 0.1% Tween20, 50 µg/mL heparin, dilution in ultrapure water) at 37ºC. Finally, organoids were washed twice for 5 minutes with 2 mL of SSCT (0.1% Tween20 in 5x SSC buffer, Sigma) at room temperature.

Amplification stage: organoids were incubated with 1 mL of amplification buffer for 10 minutes at room temperature. Meanwhile, hairpin mixtures were prepared with 100 µL of amplification buffer and 2 µL of each pair of snap-cooled hairpin probes (amplification probes, 3 µM stock). Each hybridization probe pair had two corresponding amplification probes that would bind two hybridization probe pairs. Due to their cross-reactivity (the chain reaction could be prematurely triggered if they were in contact), these hairpin amplification probes were snap cooled separately by heating them up to 95 ºC for 90 seconds and cooling them down to room temperature in a dark drawer for 30 min. Once the pre-amplification time was over, the solution was removed and 100 µL of hairpin mixture were added to the organoids, which were then incubated overnight (12-16h) in the dark at room temperature. On the next day, excess hairpins were removed with several washing steps using 2 mL of 5x SSCT at room temperature: first, 2 × 5-minute washes; then 2 × 30-min washes and finally one 5-minute wash. Samples were then mounted on µ-Slide 18 Well IBIDI chambers (81811-IBI, ibidi) and immobilized with 1% low gelling temperature agar (Merck/Sigma). When the agar solidified, 18% Optiprep (D1556-250ML, Sigma) in RNAse free PBS was added as a mounting medium and organoids were imaged in the following week. HCR probes were designed and synthesized by Molecular Instruments and their sequence is confidential.

**Table S5.** List of HCR probes from Molecular Instruments (confidential sequence).

| Hybridization Probe (target gene) | Amplification Probe (adaptor-fluorophore wavelength) |
| --- | --- |
| PTCH1 | B2-514 |
| SFRP2 | B1-488 |
| FOXA2 | B5-647 |
| FOXG1 | B4-594 |
| NKX2-1 | B4-594 |

### Imaging in situ HCR stained organoids

Neural organoids were imaged using a Zeiss LSM980 system in lambda scanning mode followed by spectral unmixing and processed with Fiji. Acquisition mode parameters were the following: pinhole size = 20 µm (increased light collection in case of very low signal), scan speed = 6, bidirectional scanning, 2x averaging per frame, averaging method = sum intensity, 8 bits per pixel. A water immersion, 10x objective was used to take Z-scans spanning each entire organoid, with a Z-step size of 10 µm. Tile scans were taken when organoid size exceeded the size of the field of view, and they were later stitched using the ZEN Blue software (v3.5.093.00008). Laser levels for each probe and fluorophore were adjusted to obtain the maximal dynamic range across all organoids, avoiding signal saturation.

### Forebrain axis scoring

To calculate a forebrain axis score per cell, forebrain progenitors (Telencephalic and hypothalamic progenitors) were subsetted from the main Seurat object, as this score would only be representative of the D-V axis at this A-P axis position. The corresponding expression matrix was extracted for these cells and for the union of reference genes involved in dorso-ventral patterning (known from the literature) with the variable features from this dataset. This expression matrix was transposed and converted into a Seurat object to observe genes as the variable of interest. ScaleData, RunPCA and FindNeighbours were run to find gene modules with correlated expression patterns. Next, we took the Nearest Neighbour graph embeddings and converted them into a symmetric matrix of neighboring (covarying) genes, which was converted into a summary data frame of correlating genes. The dorsalizing genes were separated from the ventralizing and their nearest neighbor genes were assigned to the same gene module to then calculate a dorsal and ventral score using Seurat’s function AddModuleScore. DV_score and FBaxis_score was calculated as the difference between the dorsal and ventral score, and FBaxis_rank as the cellular rankings stemming from the FBaxis_score.

### Pseudo-bulk analysis of cell line variation

To summarize gene expression per condition taking cell line and experiment information into account, the Seurat functions “AverageExpression” and “AggregateExpression” and the custom function “summarize_data_to_groups” were used. Taking the Condition_ident_line column of the metadata as grouping variable, the average normalized data matrix, the aggregated count data matrix, and the summarized metadata table were given as input to create a new Seurat object (with the “CreateSeuratObject” function). After this, the same downstream processing as earlier was followed, with SCTransform normalization and scaling, cell cycle score regression, PCA calculation with variable features from the SCT assay and UMAP dimensionality reduction.

### Morphogen-Regulon network computation

To calculate regulons, we used SCENIC ^42^ using the pyscenic implementation ^70^ (v0.12.1). First, we ran the full SCENIC algorithm with default parameters using the clustered v10 motif database ^42^. This led to the detection of 443 regulons (average size= 196 genes). Afterwards, we calculated regulon activity scores for every cell using AUCell.

We then used the GRNBoost2 algorithm (first step of SCENIC, arboreto package 0.1.6), to find links between morphogens and regulon activities. To look for time-effects and interactions, we explicitly added these terms as pseudo-morphogens. This led to 39 morphogen-regulons, linked to on average 358 regulons. We then merged the variables back to morphogens, adding edge-characteristics whether the morphogen-regulon link was detected using interactions or time-dependent effects. If a morphogen-regulon link was detected multiple times, we kept the strongest link based on weight (w).

For visualization, we also calculated correlations between morphogens and regulon activity scores (AUC), to add as additional edge characteristics. Pearson correlations were calculated for timing and concentration experiments separately. Finally, only morphogen-regulon links with a weight>200 were kept, and non-neural as well as undifferentiated cluster-associated regulons were removed from the network. The layout was computed using layout_as_backbone (keep=0.3) and plotted using igraph v.1.4.0 and tidygraph v1.2.3.

## Supplementary Files

| File name | Content |
| --- | --- |
| Sup_cluster_markers_1 | Cluster markers from neural organoid morphogen screen (Figures 1-5) |
| Sup_cluster_markers_2 | Cluster markers from neural induction method experiment (Figure 5) |
| Sup_cluster_markers_3 | Cluster markers from MiSTR vs MiSTR-like organoids experiment (Figure 6) |
| Sup_regulons_BMP4 | List of regulons associated to BMP4 treatments and corresponding details |
| Sup_regulons_BMP7 | List of regulons associated to BMP7 treatments and corresponding details |
| Sup_regulons_SHH | List of regulons associated to SHH treatments and corresponding details |
| Sup_regulons_FGF8 | List of regulons associated to FGF8 treatments and corresponding details |
| Sup_regulons_RA | List of regulons associated to Retinoic Acid treatments and corresponding details |
| Sup_regulons_CHIR | List of regulons associated to CHIR treatments and corresponding details |
| Sup_regulons_XAV939 | List of regulons associated to XAV939 treatments and corresponding details |

## Supplementary Figures

**Fig. S1.**
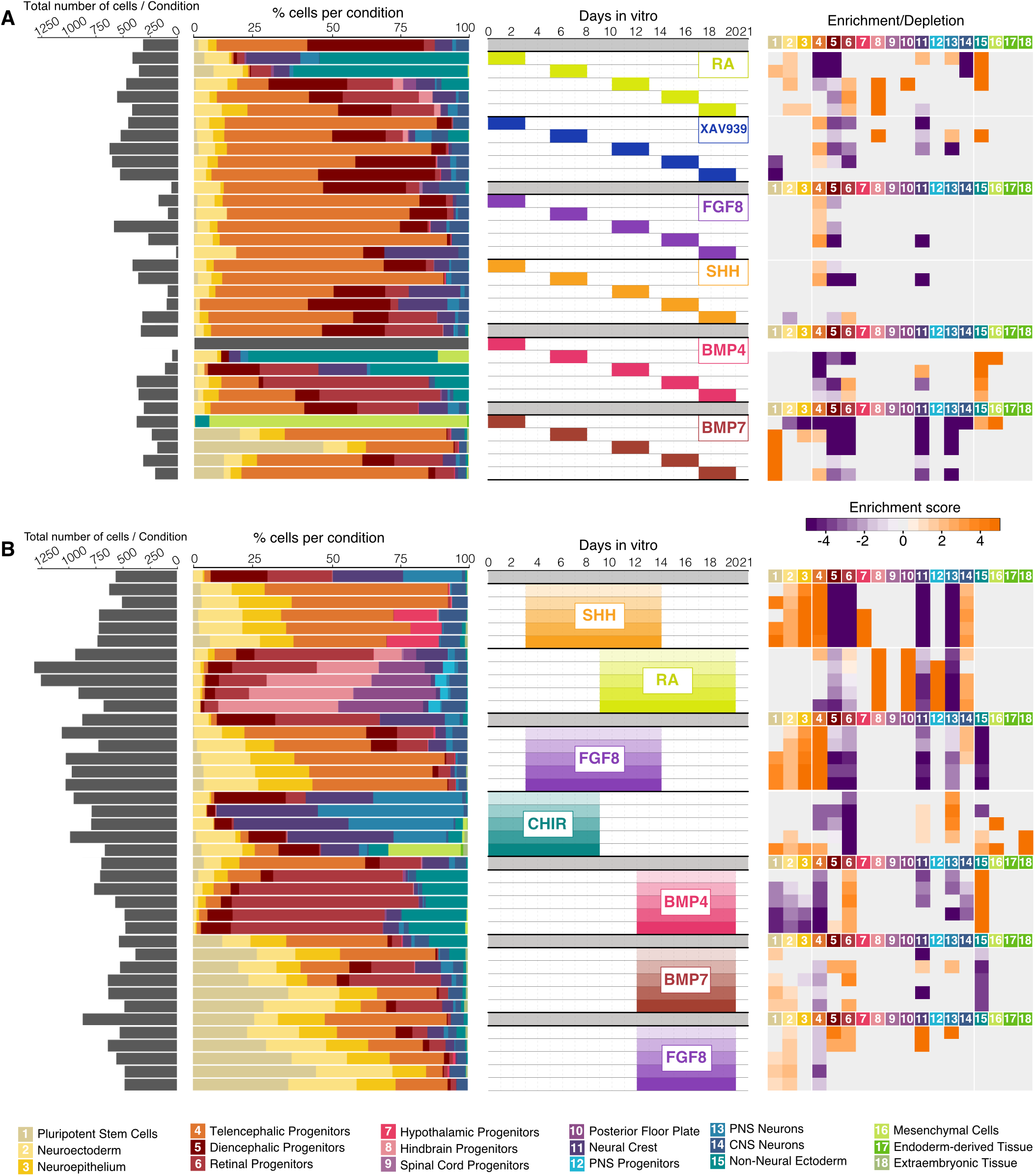
Overview of conditions and results of the timing and concentration morphogen screening (related to main figure 1). (A,B) Barplots (left) show total number of cells sequenced and percentage of cells in each cluster per condition in the morphogen exposure timing (A) and concentration (B) experiments. Heatmap shows enrichment (orange) and depletion (purple) of each cluster across conditions (right). All results correspond to the cell line HES3.

**Fig. S2.**
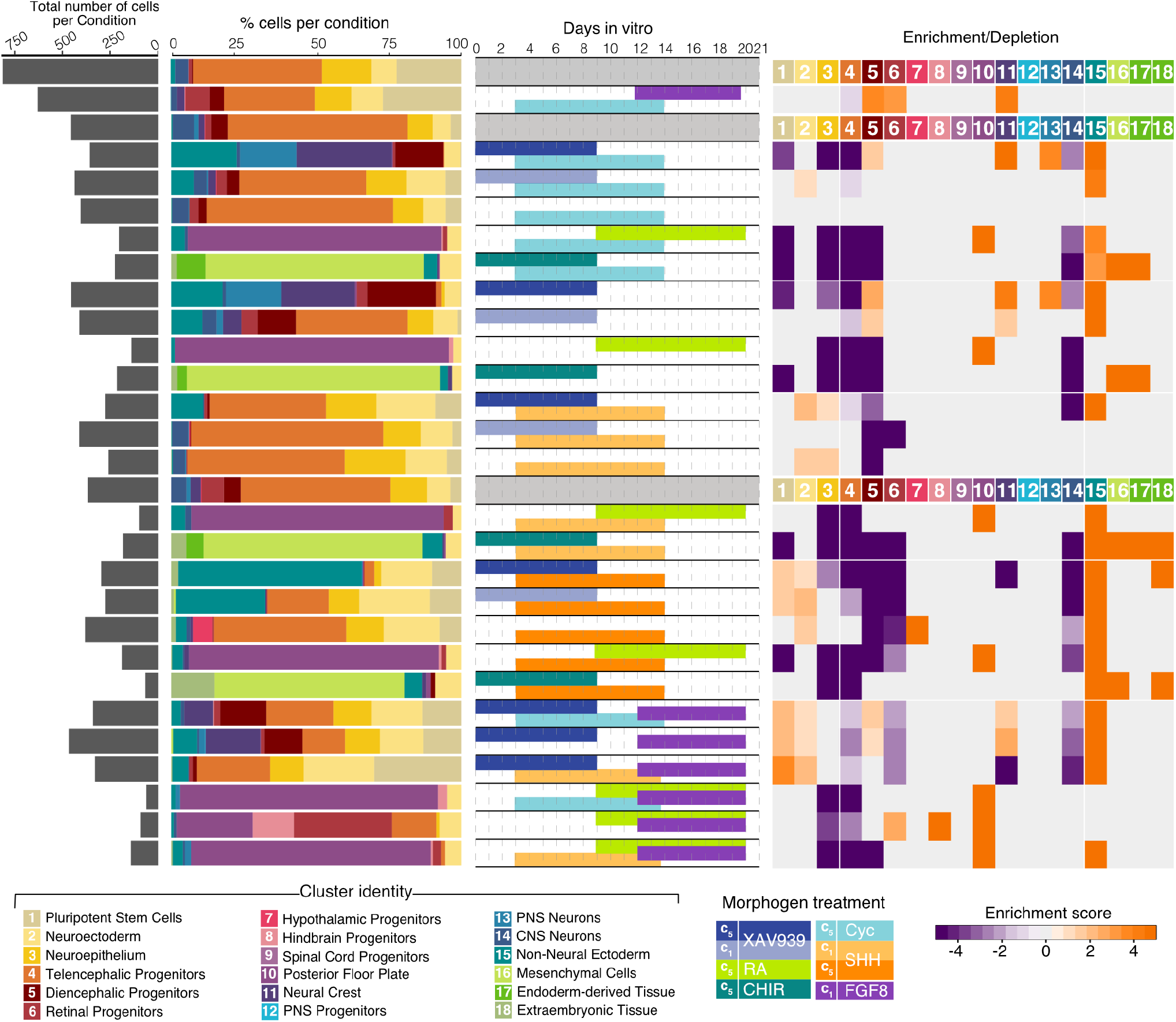
Overview of conditions and results of the combinatorial morphogen screening (related to main figure 1). From left to right, cell numbers per condition, bar plots showing cell type composition of each condition, experimental timeline for morphogen treatment, and enrichment/depletion heatmaps. All results correspond to the cell line HES3.

**Fig. S3.**
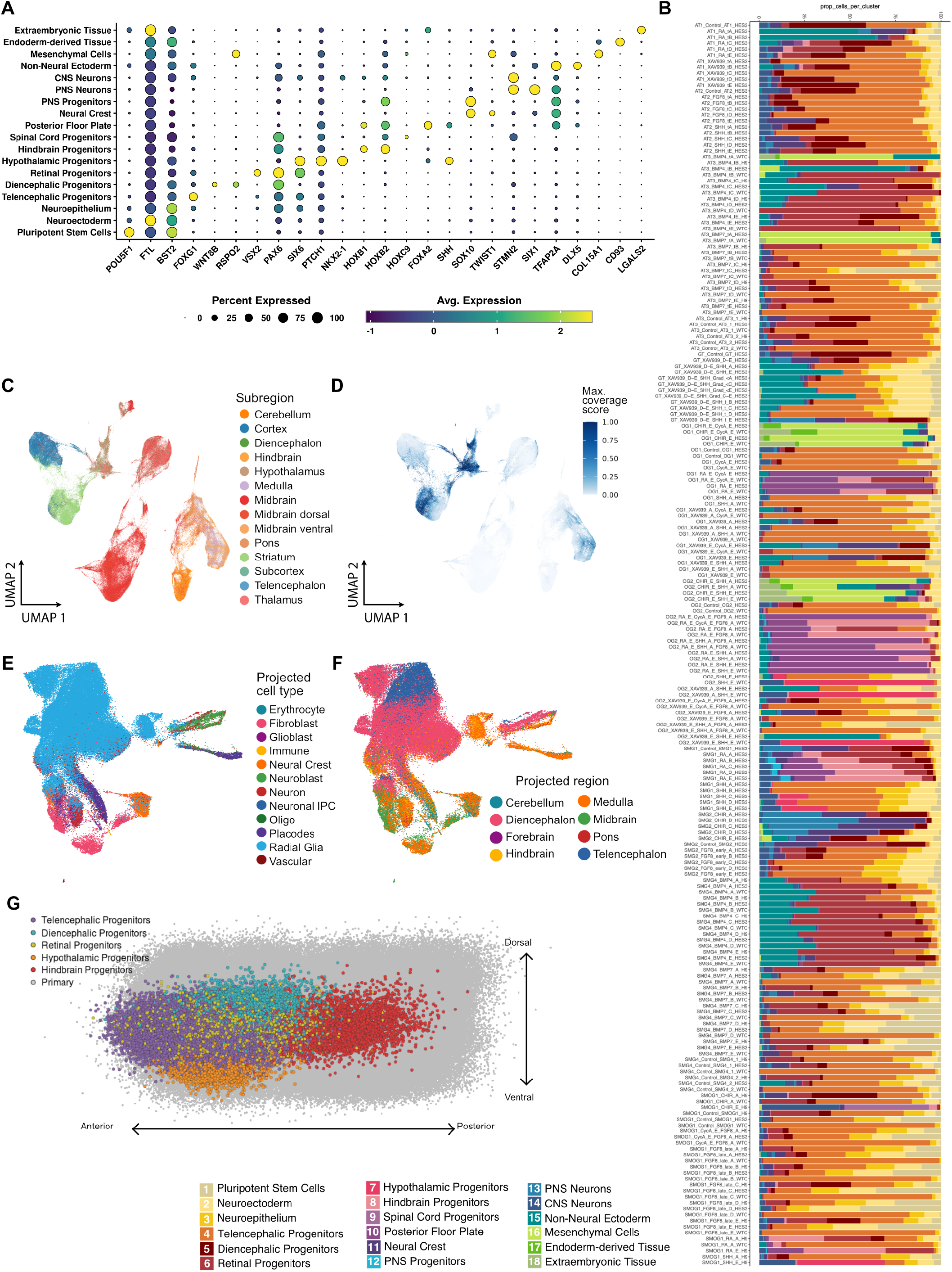
Annotation of regionalized organoid single-cell dataset and projection to existing reference datasets (related to main figure 1). (A) Dotplot showing cluster marker expression for the entire morphogen screen dataset. (B) Cell type composition barplots for all the conditions, separated by experiment and cell line. Experiment abbreviations: AT1/2/3, Timing experiments 1/2/3; GT, Gradient Timing experiments; OG1/2. Combination experiments 1/2; SMG1/2/4, Concentration experiments 1/2/4; SMOG1, mixed experiment including conditions with concentration steps and combination of morphogens. A-E correspond to concentration steps c1-c5, whereas tA-tE correspond to successive morphogen exposure windows, being tA the earliest and tE the latest (see Table S1). Cell line names are indicated at the end. (C) UMAP projection of the early radial glia from the developing human brain atlas ^26^, colored by region. (D) Same UMAP projection showing the coverage values for the morphogen patterning screen. (E,F) UMAP-RSS projection with cells colored according to the cell type (E) and region (F) they project to in the developing human brain atlas ^26^. Dataset mapping was performed using scArches ^71^. (G) Pseudo-coordinate system embedding showing the spatial mapping of neural organoid regional cell types to different locations along the dorso-ventral and rostro-caudal axis of the developing human brain between weeks 5 and 7^26^. Gray dots represent the cells from primary tissue spread out according to their dorso-ventral and rostro-caudal axis score.

**Fig. S4.**
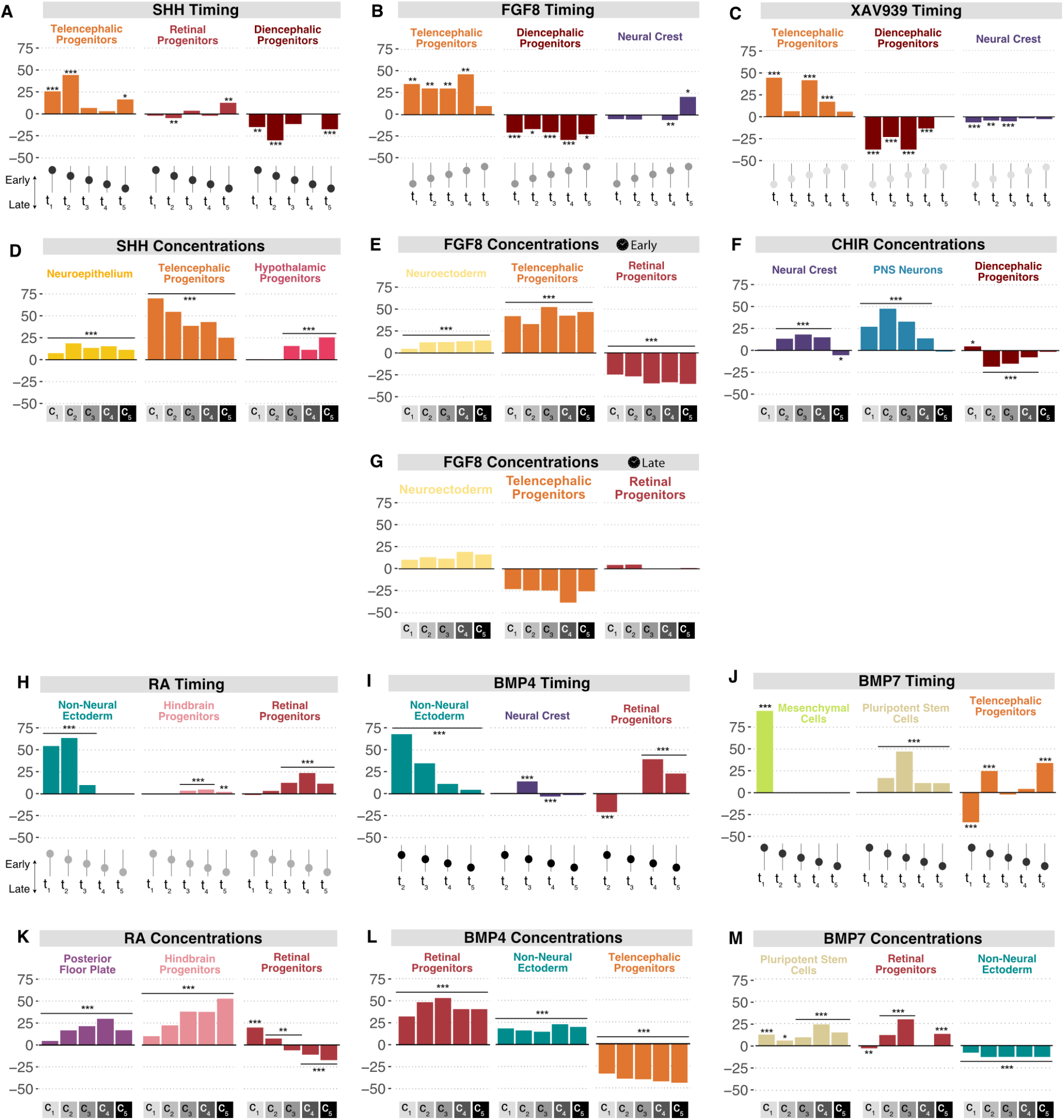
Regulation of neural and non-neural cell fates by morphogen timing and morphogen concentrations (related to main figure 2). Most significantly enriched or depleted clusters for each morphogen after single time point pulses (Timing) or prolonged exposure to concentration steps (Concentrations). *** p-value < 0.001, ** 0.001< p-value < 0.01, * 0.01 < p-value < 0.05 (Bonferroni adjusted).

**Fig. S5.**
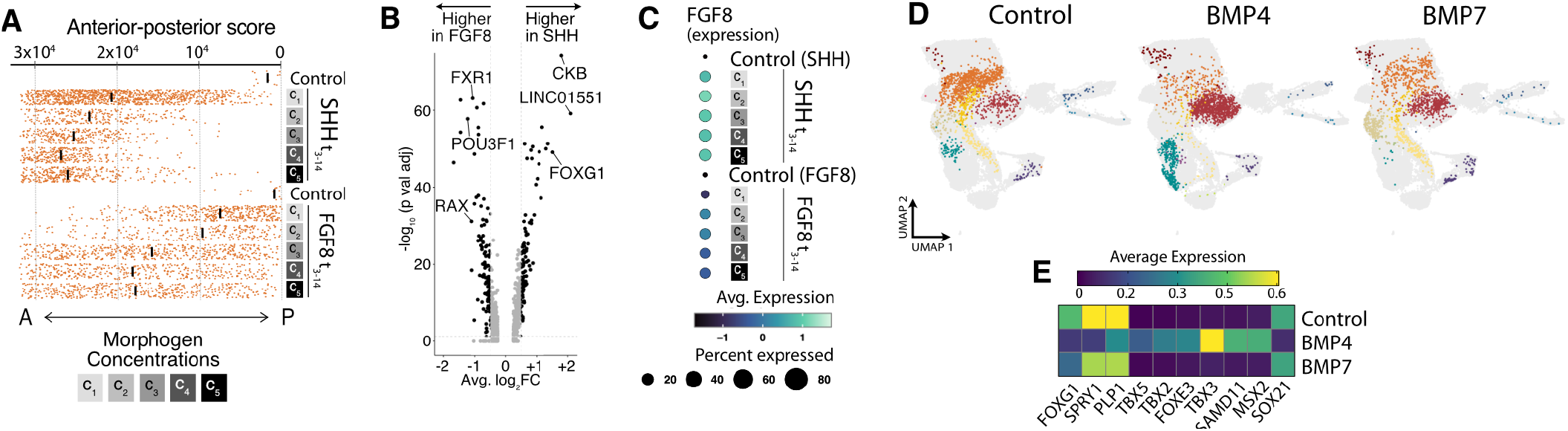
Comparing the effects of FGF8 versus SHH and BMP4 versus BMP7 treatment (related to main figure 2). (A) Antero-posterior scores for telencephalic progenitors treated with SHH or FGF8 concentration steps. (B) Volcano Plot showing the differentially expressed genes between the FGF8 c3 t3-14 and SHH c2 t3-14 induced telencephalic progenitors. (C) Endogenous FGF8 expression in telencephalic progenitors in response to SHH or FGF8 concentration steps applied at the same time window. SHH expression not shown, as it had consistent zero values. (D) UMAP representation of cell fate-shifting under BMP4 and BMP7 treatment in comparison to control. Only cells from the BMP concentration steps experiment are shown and are coloured by cluster. (E) Heatmap showing differentially expressed genes in ectodermal cells from control, BMP4 or BMP7 conditions. *** p-value < 0.001, ** 0.001< p-value < 0.01, * 0.01 < p-value < 0.05 (Bonferroni adjusted).

**Fig. S6.**
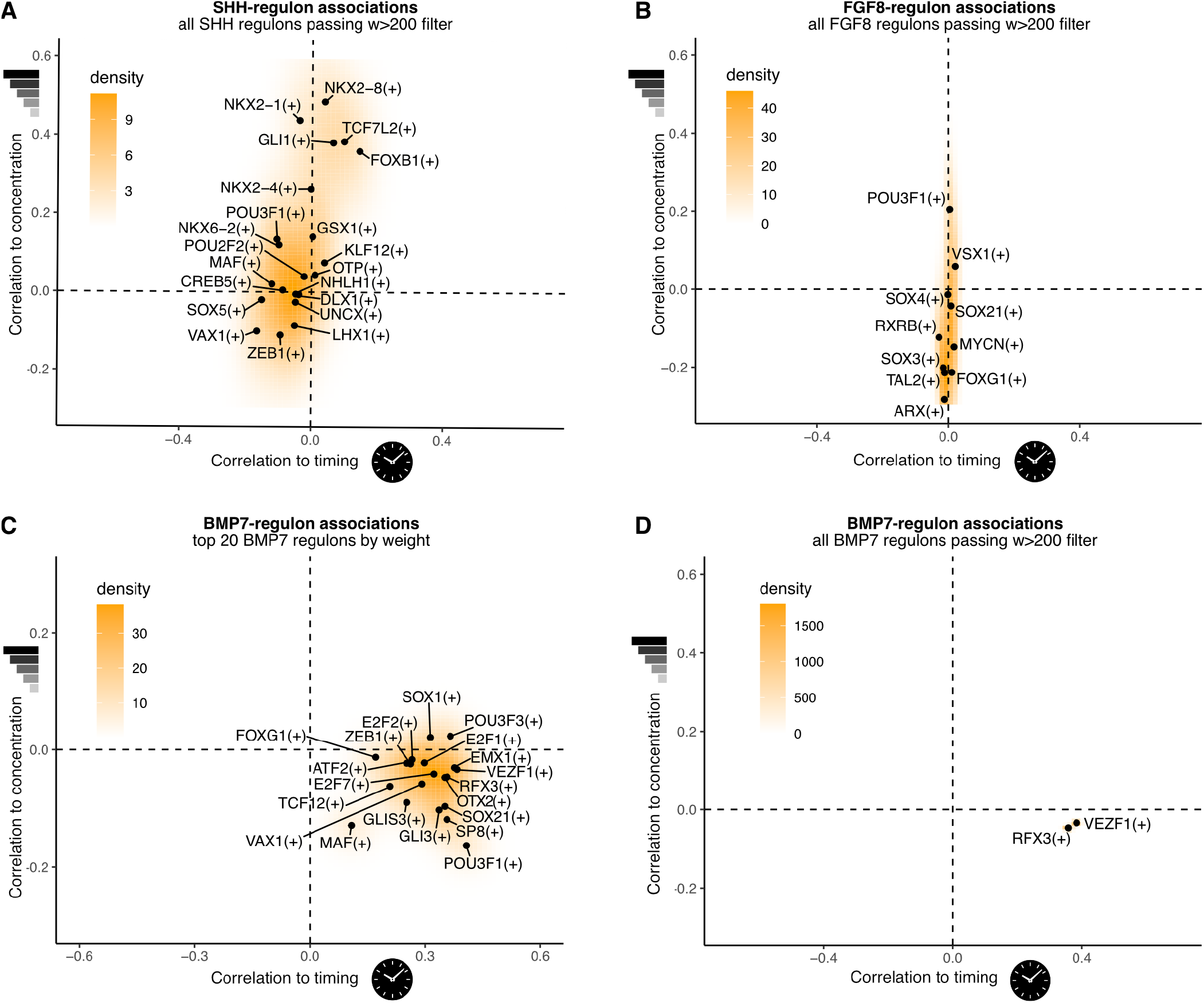
Relationship between timing and concentration on regulon activity (related to main figure 3). Scatterplot shows the relationship between correlation to timing (x-axis) and concentration (y-axis) of morphogen associated regulons, with selected regulons labeled. (A) Scatterplot of SHH-associated regulon correlation to timing and concentration, showing wide dependency on both variables. (B) Scatterplot of FGF8-associated regulon correlation to timing and concentration, showing wide dependency on morphogen concentration. (C,D) Scatterplot of BMP7-associated regulon correlation to timing and concentration, showing roughly equal dependency on both variables. (C) shows the top 20 weighted regulons for BMP7, while (D) shows only the ones above a weight value of 200.

**Fig. S7.**
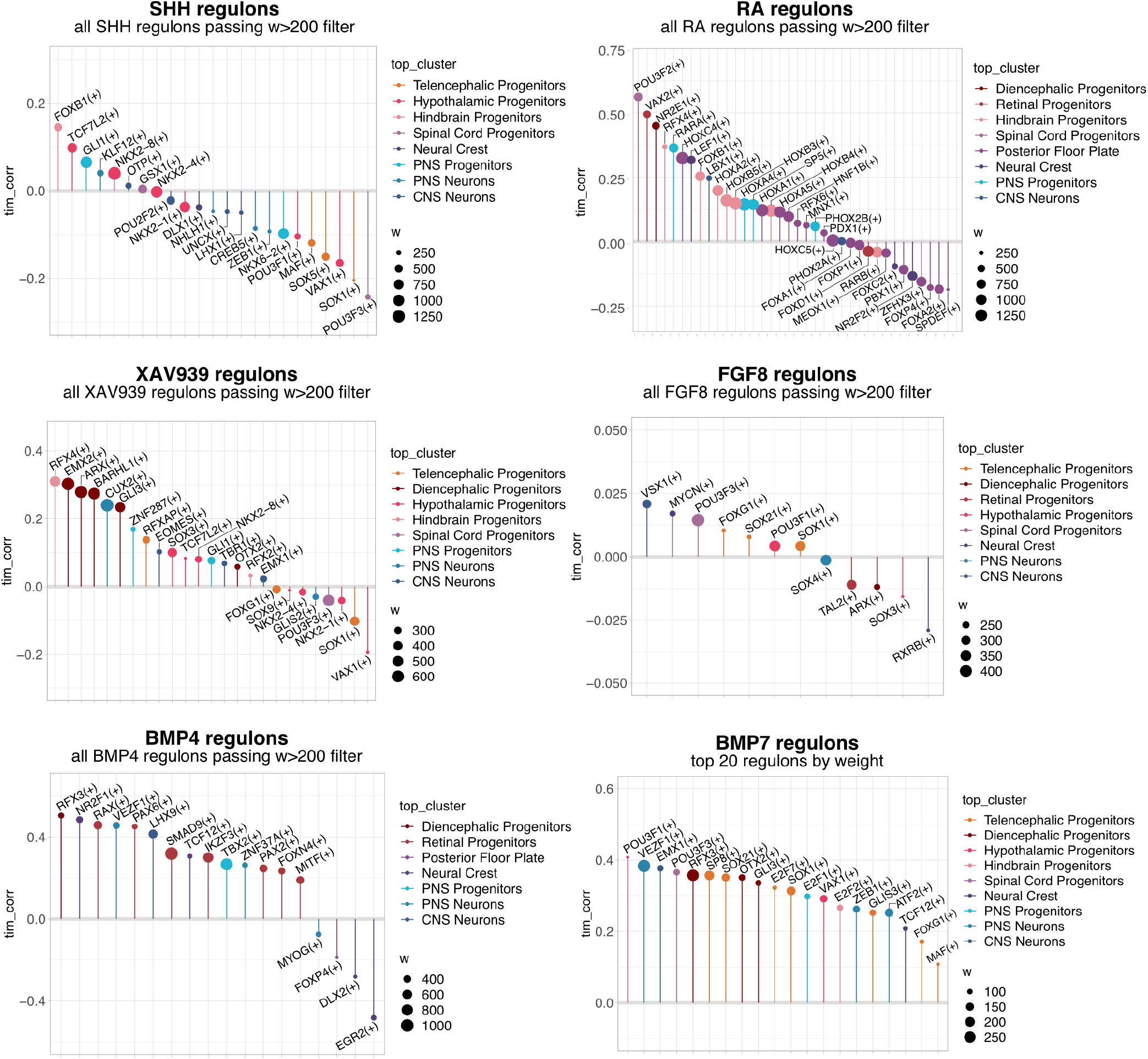
Treatment timing correlation of morphogen-associated regulons (related to main figure 3). Lollipop plots showing the correlation to treatment timing from each of the regulons associated to each morphogen. The size of the dot on top represents the weight of the corresponding association to the morphogen, and the color represents the cluster where the regulon is enriched. For all morphogens except BMP7, the top regulons with associations to timing surpassing weights of 200 are shown. In the case of BMP7, as only VEZF1(+) and RFX3(+) showed weights over 200, the top 20 regulons scoring highest correlation to timing are shown.

**Fig. S8.**
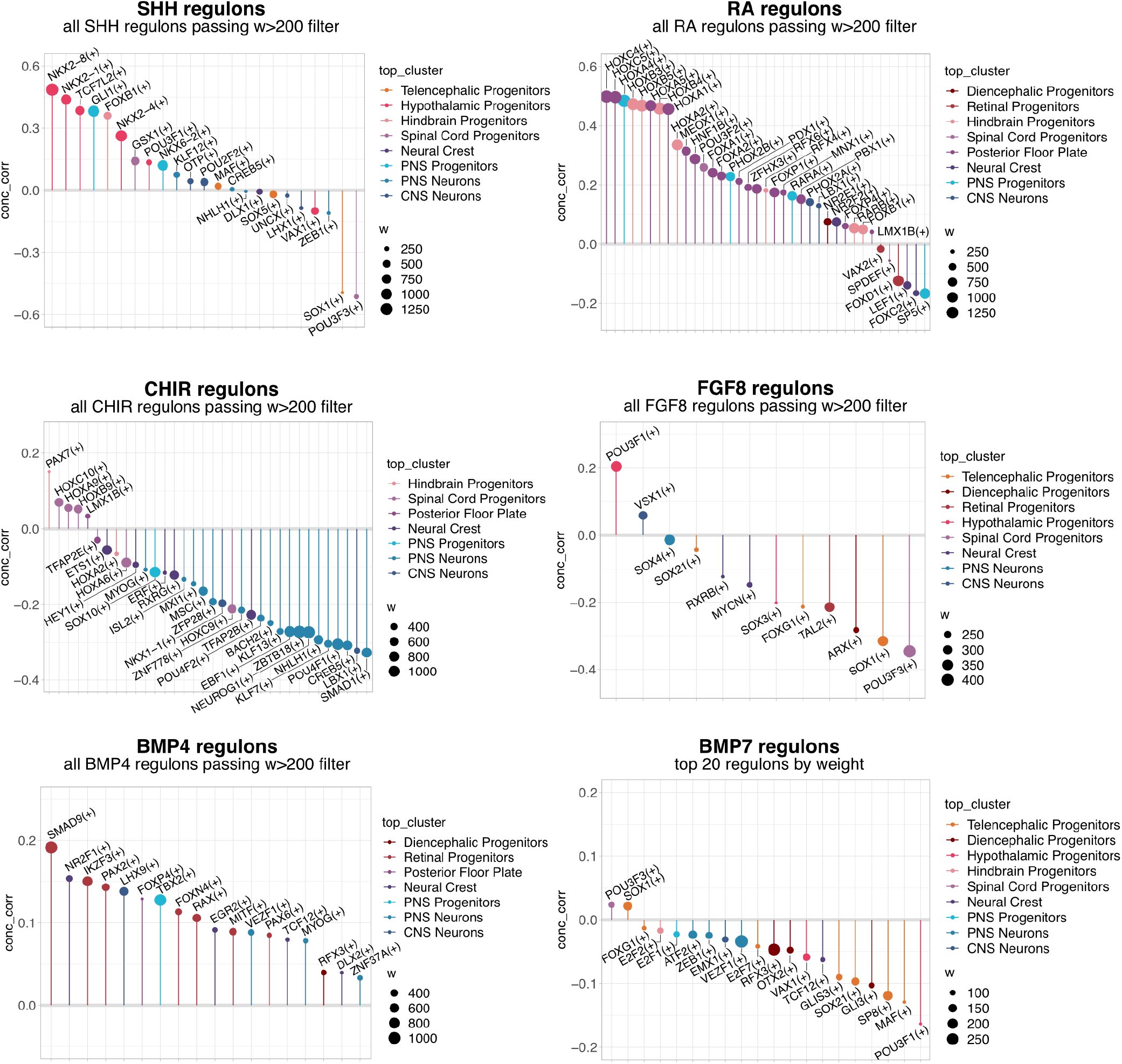
Treatment dose correlation of morphogen-associated regulons (related to main figure 3). Lollipop plots showing the correlation to treatment dose from each of the regulons associated to each morphogen. The size of the dot on top represents the weight of the corresponding association to the morphogen, and the color represents the cluster where the regulon is enriched. For all morphogens except BMP7, the top regulons with associations to concentration surpassing weights of 200 are shown. In the case of BMP7, as only VEZF1(+) and RFX3(+) showed weights over 200, the top 20 regulons scoring highest correlation to concentration are shown.

**Fig. S9.**
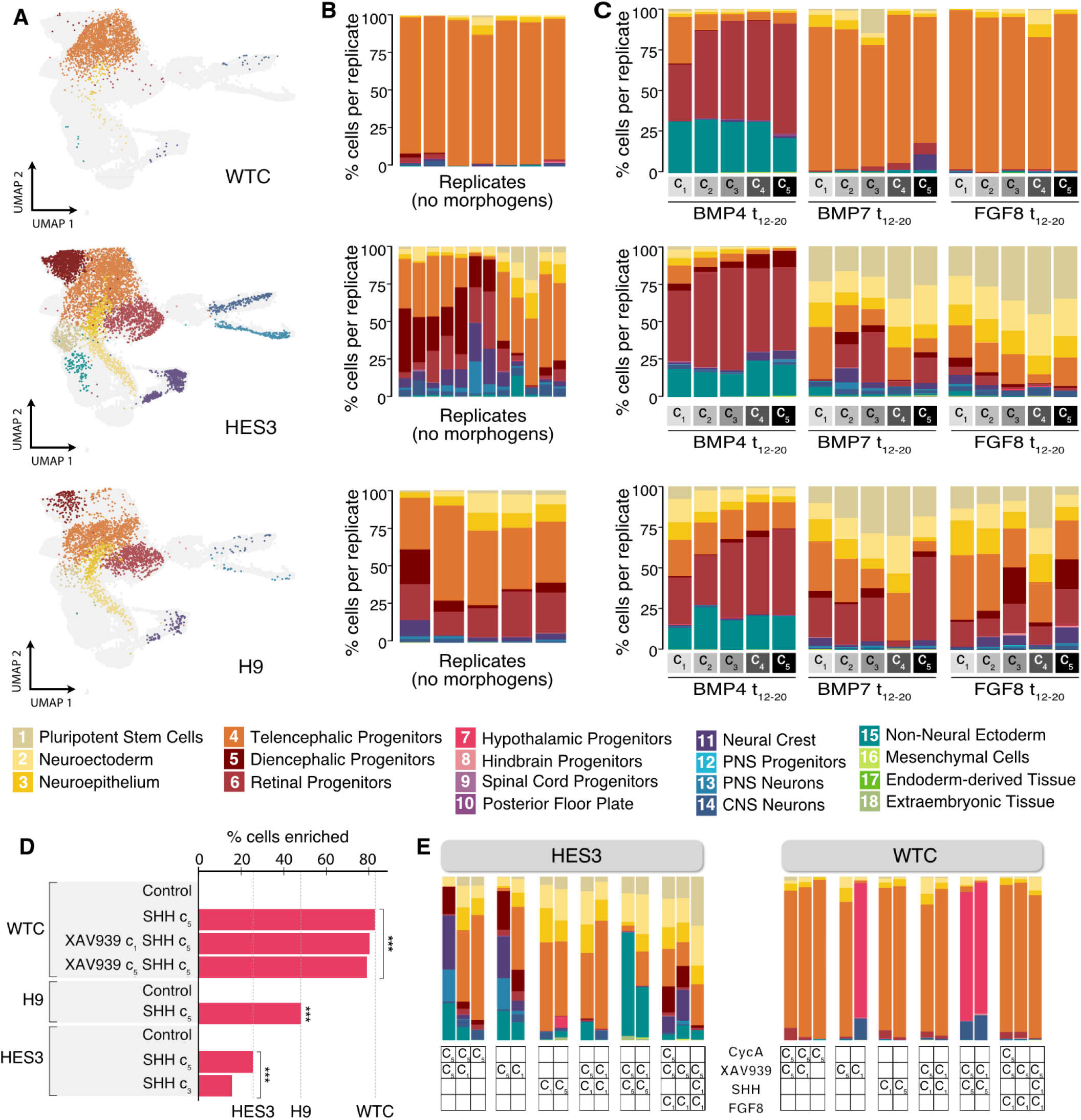
Morphogen-induced patterning does not completely override hPSC line regional biases (related to main figure 5). (A) UMAP projections from figure 1 showing the clusters produced by control organoids of each cell line. (B) Corresponding composition bar plots for each cell line replicate. (C) Cell type composition of neural organoids exposed to increasing concentration steps of BMP4, BMP7 and FGF8 from day 12 to 21. Top, WTC organoids, middle, HES3 organoids, bottom, H9 organoids. (D) Enrichment levels of hypothalamic progenitors achieved with SHH-based treatments in three different hPSC lines. (E) Comparison of HES3 and WTC responses to combinatorial patterning with XAV939, Cyclopamine, SHH and FGF8 at the cell type level, showing the influence of each cell line’s regional tendencies on the generated cell types. The concentration steps used for each morphogen are indicated in each cell of the bottom grid (c1/c5). Empty cells indicate that the particular morphogen was not included in that condition. *** adjusted p-value < 0.001.

**Fig. S10.**
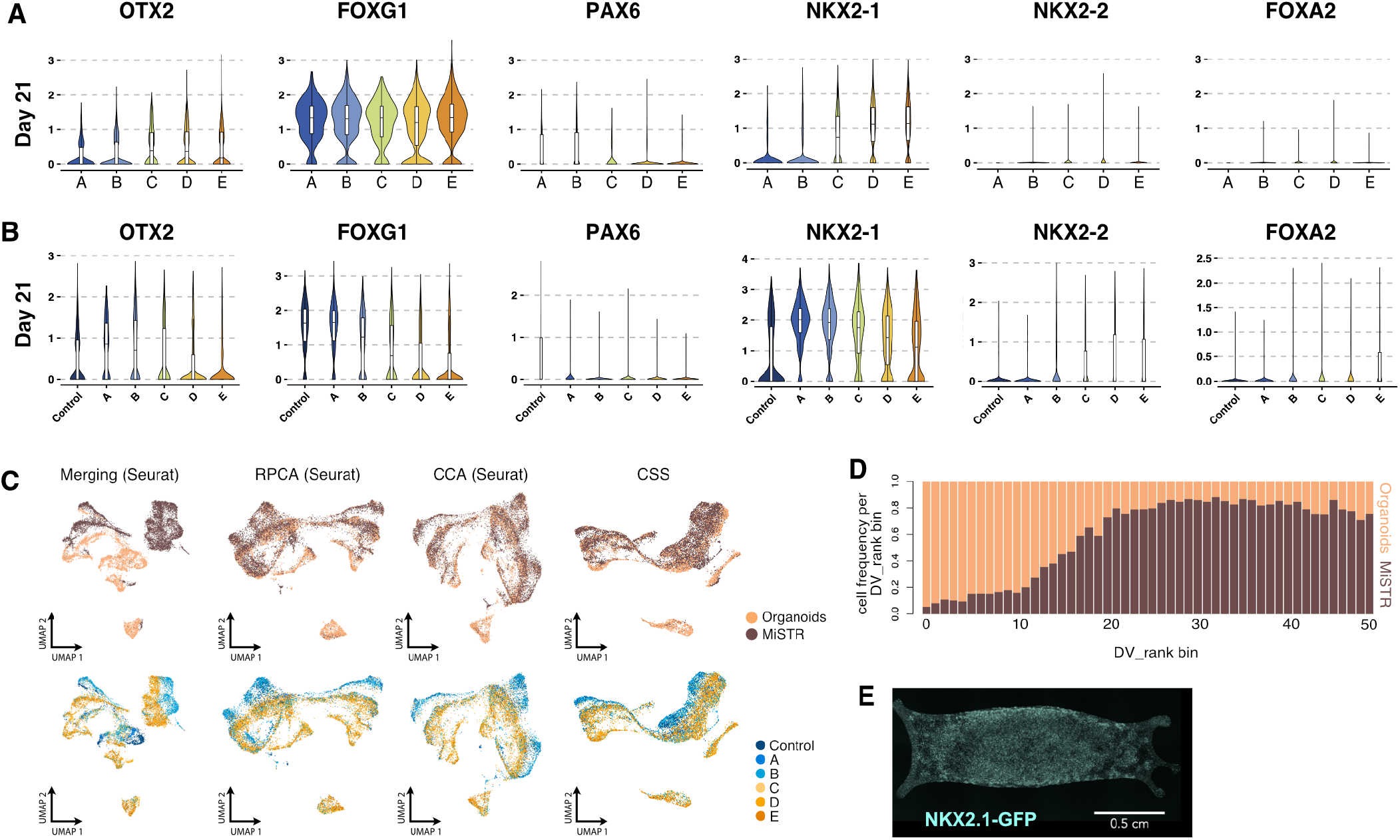
Characterization and data integration of MiSTR and MiSTR-like organoid cultures (related to figure 6). (A,B) Expression patterns of key forebrain patterning genes across sections A-E in MiSTR tissue (A) and MiSTR-like organoids (B), showing a tendency towards higher ventralization in organoids. (C) UMAP resulting from different methods showing the robustness of the integration. (D) Distribution of MiSTR (brown) and MiSTR-like organoid cells (orange) across the binned DV ranks, showing coverage of the same dorso-ventral domains at different frequencies. (E) Fluorescence image showing widespread NKX2.1-GFP expression in condition “D” MiSTR tissue cultured in the absence of PDMS microfluidic tree. The image was taken on differentiation day 12.

## Notes

### Competing Interest Statement

The authors have declared no competing interest.

